# A minimal visual world model predicts exploration of naturalistic landscapes in flies

**DOI:** 10.64898/2026.03.04.709519

**Authors:** Thomas F. Mathejczyk, Gerit A. Linneweber

**Affiliations:** Institut für Biologie Abteilung Neurobiologie Fachbereich Biologie, Chemie und Pharmazie Freie Universität Berlin

## Abstract

Animals, including small insects, can explore complex natural environments to locate resources and potential mates. How a miniaturized nervous system extracts actionable structure from complex visual scenes to guide effective exploration remains incompletely understood.

To bridge the gap between behavioristic experimental control and ethological realism, we combined a virtual-reality flight assay with naturalistic three-dimensional environments based on high-resolution satellite imagery of real-world landscapes. Flies that had never experienced natural environments exhibited individually stable, non-random exploration patterns and preferred flying over vegetated and elevated terrain while avoiding water-like features, indicative of a minimalistic world model for ecologically relevant terrain selection. Flight paths were characterized by saccadic turns, which we analyzed in relation to visual scene components, including brightness, color composition, optic flow, elevation, and contrast. Using multivariable regression modeling, we identified a hierarchy of visual cues influencing saccade frequency, amplitude, and directionality, and mapped the retinal areas most strongly gating saccadic decision-making. We discovered individual differences in motion-sensitivity thresholds that predict stable individual exploration-exploitation phenotypes. indicating a population-level bet-hedging strategy to increase terrain coverage and resource encounter rates. Individuality-based probabilistic decision-making simulations, utilizing our hierarchical visual models, successfully replicated exploration behavior, with the predicted dispersal and trajectory statistics matching the empirical data and demonstrating generality even for visual assays not included in the training dataset. Our work links individual visual decision rules to efficient exploration mechanisms of large-scale natural landscapes, potentially inspiring neural circuit theory, ecological prediction frameworks, vision-based pest control, and energy-efficient autonomous system design.

## Introduction

Animals constantly sample their visual environments and must select behaviorally relevant signals while filtering irrelevant sensory input. While the visual world surrounding animals in nature can be very complex, most laboratory research on vision utilizes well-quantifiable but relatively simple visual stimuli, including stationary or moving stripe patterns (Linneweber et al., 2020; Maimon et al., 2010; Mongeau & Frye, 2017), looming stimuli (von Reyn et al., 2017), white noise (Aptekar et al., 2015; Theobald et al., 2010), simple geometric patterns (Neuser et al., 2008; Reiser & Dickinson, 2010; Tang et al., 2004), simulated celestial bodies (Dacke et al., 2019), polarized light cues (Stephens et al., 1953; Warren et al., 2018), or color choice assays (Gao et al., 2008; Schnaitmann et al., 2013; Yamaguchi et al., 2010). Only recently have investigators begun to explore visually guided behaviors in response to more complex visual scenes in vertebrates (Isaacson et al., 2025; Pinke et al., 2023) and invertebrates (Haberkern et al., 2022). To bridge the gap between simplified laboratory stimuli and natural scenes, we generated a multiparametric three-dimensional (3D) virtual reality derived from high-resolution satellite data, which we refer to as *naturalistic* because it captures key properties of natural scenes under controlled laboratory conditions. Our setup consists of a tethered flight apparatus surrounded by a panoramic LED matrix that displays the simulated naturalistic environment in a closed loop, driven by the fly’s heading decisions. This allows quantitative testing of large-scale exploration behavior within naturalistic landscapes under tightly controlled laboratory conditions.

### Visually Guided Large-Scale Dispersal in *Drosophila melanogaster*

The vinegar fly, *Drosophila melanogaster,* is highly effective at extracting behaviorally relevant visual information from its surroundings, enabling it to explore large-scale natural environments. Without wind, flies can migrate in arid environments up to 15km per day (Coyne et al., 1982; Jones et al., 1981), and over several days, they can migrate even up to 48km without food (Hocking, 1953). During long-range dispersal, flies can maintain stable headings using the sun as an orientational cue, either by extracting celestial color/intensity gradients (Giraldo et al., 2018; Pae et al., 2024; Warren et al., 2019) or by using skylight polarization information (Warren et al., 2019). The use of these celestial cues allows flies to keep their direction even in the absence of any other visual landmarks (menotaxis) (Leitch et al., 2021).

### Individuality in sensorimotor processing

Even genetically similar flies reared under experimentally controlled conditions can exhibit stable inter-individual behavioral differences, including visual choice tendencies and orientation biases (Linneweber et al., 2020). Near-isogenic *Drosophila* populations show individually stable differences in phototactic preference that can be modified by neuromodulatory mechanisms (Kain et al., 2012). Similarly, spontaneous locomotive handedness biases are individually stable and are controlled by specific neurons in the central complex, indicating a circuit basis for behavioral idiosyncrasy (Buchanan et al., 2015). Importantly, it was shown that the tendency for inter-individual variability itself has heritable components, implying that natural selection can shape not only the means of behavioral traits but also the structure of behavioral variance (Ayroles et al., 2015). Therefore, we hypothesized that individually stable differences in visual processing may also lead to distinct exploratory phenotypes, shaping the exploration–exploitation balance at the population level.

### Summary and key questions

Animals routinely explore large, visually complex environments to locate sparse resources. How a small nervous system extracts actionable structure from natural scenes and converts it into efficient exploration strategies remains a central open question in the neurosciences and ecology. Here, we combined a closed-loop virtual flight simulator with a novel framework for rendering 3D, naturalistic landscapes to quantify visually guided exploration behavior in *Drosophila*. We reveal that large-scale exploration emerges from structured, cue-dependent saccadic decision-making rather than random movement. Additionally, we show that stable inter-individual differences in visual motion sensitivity give rise to distinct individual exploration–exploitation phenotypes. A compact probabilistic model trained on individual fly behavior recapitulates empirical trajectory statistics, generalizes to classical visual assays, and predicts large-scale dispersal patterns in novel environments.

Together, our work links low-level visual processing rules to emergent exploration strategies in complex landscapes, providing a mechanistic framework for understanding how small brains achieve efficient, scalable exploration and how individual variability can enhance population-level performance.

## Materials and Methods

### Animals

Experiments were performed with 4– to 7-day-old wild-type *Drosophila melanogaster* (CantonS, source: Bassem Hassan lab). Flies were reared at 25°C, 50% relative humidity and a 12/12h light/dark regimen on standard *Drosophila* yeast-cornmeal medium (64g cornmeal, 7.5g agar, 160g yeast, 85,5ml sugar cane syrup in 1l of water).

### Experimental procedure

Flies were cold-immobilized on a 4°C Peltier element (for detailed tethering station description, see: (Mathejczyk & Wernet, 2020)) and a 10×0.1mm steel pin (entosphinx.cz) was attached to the dorsal anterior thorax at an angle of approximately 60° from horizontal using UV-cured glue (Bondic). After tethering, flies were allowed to recover for at least 30 minutes before being tested in the flight simulator for 15 minutes. For multi-day individual experiments, flies were carefully separated from the steel pin and put into single vials containing food for ∼24h.

### Behavioral assay

We tested flies in a virtual flight simulator described in detail in: (Mathejczyk et al., 2024). In short, the assay consists of a magneto-tether apparatus (based on: (Bender & Dickinson, 2006)) surrounded by a cylindrical low-latency LED-based display system for presenting visual stimuli to the flies. The magneto-tether is constructed using two vertically aligned magnets. A tethered fly is placed between the magnets with the tip of the steel pin inserted into a 1mm-diameter V-shaped sapphire bearing attached to the top magnet to reduce friction. The magnetic field keeps the fly stationary within the apparatus while still allowing it to rotate freely around its yaw axis. The fly is illuminated via 850nm LEDs and filmed through the bottom ring magnet from below using a near-infrared camera (FLIR Firefly S FFY-U3-04S2M-C) at 90Hz (see: (Mathejczyk et al., 2024)). Visual stimuli were presented using a cylindrical 256×128 pixel RGB LED matrix (1.4° per pixel, 120Hz frame rate, 5kHz refresh rate) controlled via a low-latency LED controller (Novastar mctrl 660pro, 120Hz, <1ms pixel update latency) connected to the computer via HDMI (for stimuli description and spatial extent of the stimulus, see Fig. 1b). An aquarium pump was connected to the system to deliver air puffs through the bottom ring magnet of the magneto-tether to the flies to reinitiate flight if they stopped flying.

**Fig. 1:**
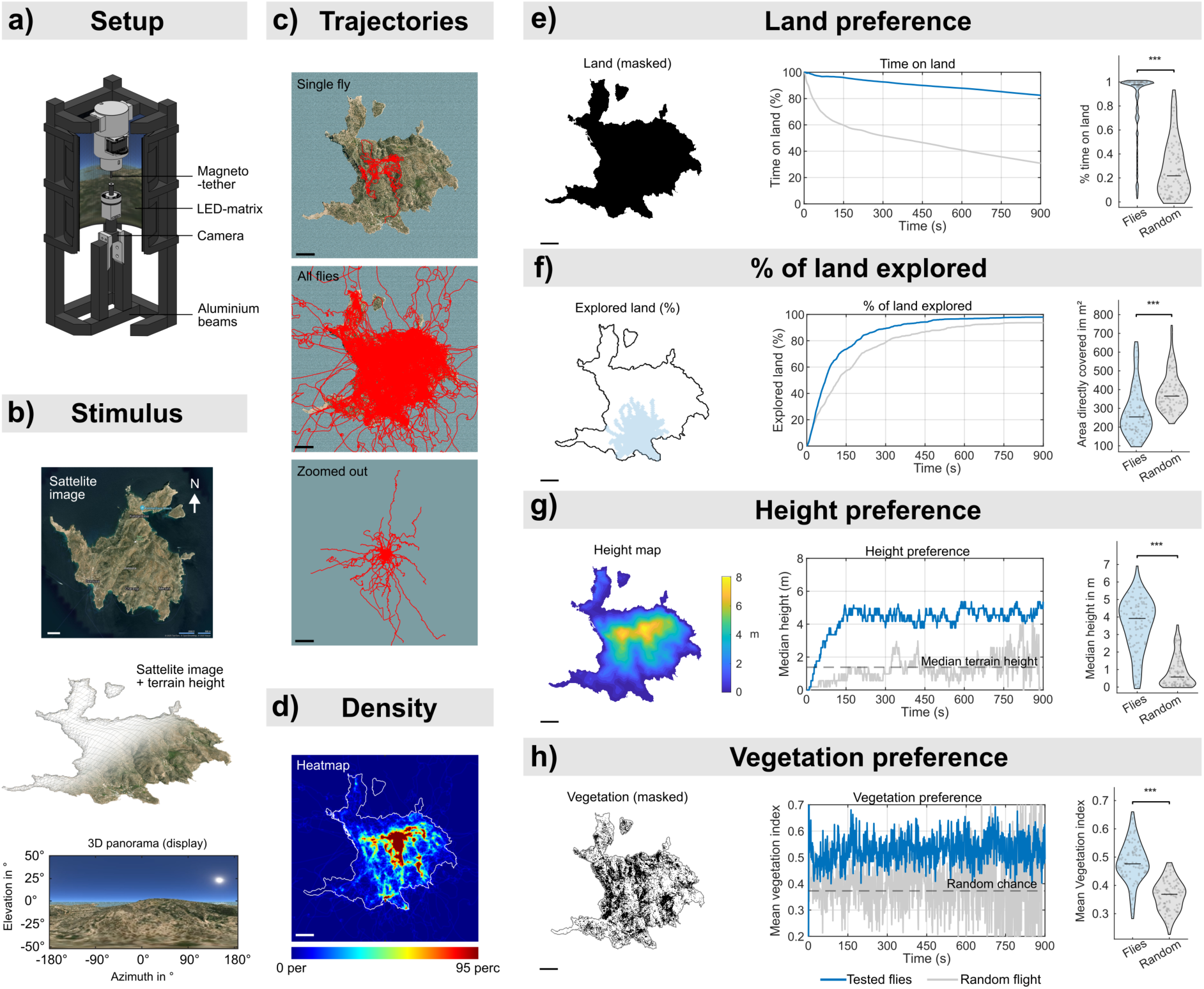
Flies explore virtual naturalistic environments with visuomotor biases towards vegetation and elevated height and away from water. a) Overview of the setup. A fly is placed within a magneto-tether and surrounded by a panoramic RGB LED matrix displaying a naturalistic virtual landscape. The fly’s heading data (tracked at 80Hz) drives a panoramic virtual camera through the 3D landscape, maintaining a constant forward velocity at a constant 2m above the ground. The panoramic camera’s output is rendered onto the LED matrix, providing closed-loop visual feedback to the flying fly. b) Stimulus design. Combining satellite image data of the Greek island Donousa (top panel) with SRTM elevation information (middle panel) to create a high-resolution 3D model of the island. Bottom panel: rendered panorama as displayed on the LED matrix. A sky and a sun texture, as well as animated reflective water, were added to the visual scene. **c)** Overview of 2D trajectories of the tested flies. Top panel: representative fly, middle panel: all tested flies (n=106, 53 flies tested on 2 consecutive days), bottom panel: zoomed-out middle panel. When flies fly over water, they fly rather straight paths at a constant but arbitrary angle relative to the sun. Scale bars: top 20m, middle 20m, bottom 200m. **d)** Heat map of all fly trajectories (n=106). Flies explore the entire island and show elevated detection probabilities near the hills (see the height map in Fig. 1g). The color scale was cropped to the 95th percentile of the density data for better visibility. Scale bar: 20m. **e-f)** Flies show innate visual preferences that increase evolutionary fitness in natural environments. Blue traces show preferences of tested flies (n=106) and grey traces represent simulated flies performing random flight (n=106). The test window was 15min. Scale bars = 20m. The data was analyzed using the Wilcoxon rank sum test with asterisks indicating p-values: *p<0.05, **p<0.01, ***p<0.001. **e)** Flies prefer flying over land. Left: masked island. Middle: percentage of flies flying over land. Right: vase plots showing the individual percentage of time each fly spent flying over land. **f)** Flies explore almost the whole island. Left: explored area after 30s (blue, 2m exploration radius around each fly). Middle: total percentage of the island explored. Right: vase plots showing the area explored by each fly. **g)** Flies prefer flying over high elevations. Left: height map of the island. Middle: median height of the fly population. Right: vase plots showing the mean individually preferred elevation on the island. **h)** Flies prefer flying over vegetation. Left: masked vegetation (black) on the island. Middle: mean vegetation preference of the fly population. Right: vase plots showing the mean individual vegetation preference when flying over land.

### Closed-loop tracking and 3D virtual reality framework

Flight heading was extracted using a custom MATLAB script (see Extended Data Fig. 1a) at ∼80Hz. We applied a virtual forward velocity of 2m/s to the extracted fly headings to compute 2D trajectories whose coordinates were sent to an instance of the Blender game engine (Blender Foundation, version 2.79b) via UDP connection. The 2D coordinates were used to translate a panoramic camera (2m above the ground) through a 3D naturalistic environment, rendering a 256×128-pixel image displayed on the LED matrix surrounding the fly. As a naturalistic environment, we chose the Greek island Donousa, which was digitized using the Blender GIS addon to combine satellite image data (https://www.bing.com/maps) with elevation data (SRTM data, 30m resolution). We chose the island since it is small, only sparsely populated, roughly circular, and offers high-resolution satellite data. The island was scaled down to approximately 150mx150mx8m (WxLxH). We rotated the island so that Stavros Beach (starting position for flies) was centered in the virtual south (see Fig. 1b for original orientation). Throughout the study, we refer to the cardinal directions based on this rotated version. A sky texture with a sun (southeast) was added to the scene, along with an animated, reflective water texture surrounding the island (see Fig. 1b, bottom panel). All flies started at the same point on Stavros Beach and were free to explore all regions of the island/water. To ensure similar start headings across experiments, flies were presented with a static image of a thin (∼5° horizontal extent) vertical black bar on a white background, positioned in the virtual north. When a fly maintained a heading stable around ±5° from the bar’s center for 1s, the panoramic camera output from Blender was displayed, and the experiment began.

### Data analysis

If not stated otherwise, all data analysis was done using MATLAB. After the experiments, 2D trajectories were resampled to 80Hz using interpolation for even per-frame step lengths. Trajectory data 5s before and after each landing was excluded from analysis. An LLM (ChatGPT) was used for coding assistance. All code was manually reviewed for correctness and reliability of data output.

### Saccade detection

Saccade detection was based on the detailed description in: (Mongeau & Frye, 2017). In short, the angular velocity during flight was calculated per frame for each fly from heading data and then smoothed using the peak-preserving Savitzky-Golay filter in MATLAB. A noise threshold was calculated from each fly’s median absolute angular velocity divided by 0.6745. Whenever the angular velocity peaked above this threshold, the rotational bout was quantified as a saccade (see Fig. 2a). Saccade amplitude was defined as the angular difference in flight heading between saccade start and end.

**Fig. 2:**
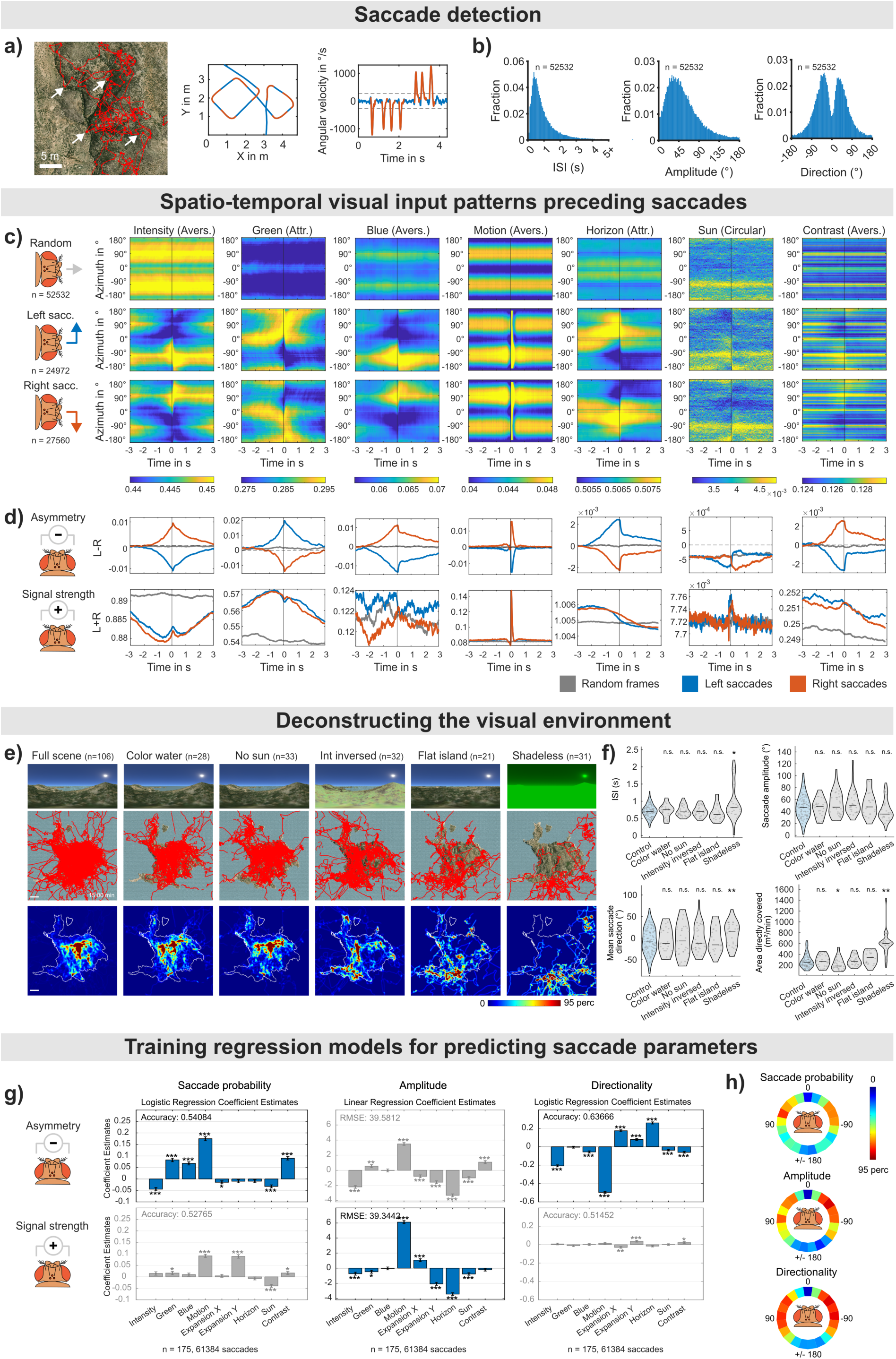
Visually guided saccade models explain fly flight behavior. **a)** Saccade detection (based on: (Mongeau & Frye, 2017)). Left: zoomed 2D trajectory of a single fly showing rapid changes of heading (saccades, exemplary white arrows) interspersed by segments of straight flight (inter-saccade ntervals, ISI). Scale bar = 5m. Middle: section of the trajectory on the left (red: saccades, blue: ISI). Right: angular velocity of the trajectory shown in the middle panel over time. For each fly, a noise threshold was calculated (Grey dotted lines). Whenever the angular velocity crossed this threshold, the rotational bout was quantified as a saccade (red). **b)** Resulting saccade parameters when flying over land (n=106, 52532 saccades quantified in total). Left: histogram of ISI. Middle: histogram of absolute saccade amplitudes. Right: histogram of saccade directions. **c-d)** Spatio-temporal visual input patterns influence saccade responses. **c)** averaged spatio-temporal azimuthal visual input patterns of non-saccadic periods and all detected saccades aligned at time point 0s (land-based saccades only). Top row: average visual input 3s before/after randomly sampled, non-saccadic frames for seven of the quantified visual parameters (n=106, 52532 saccades). Middle row: average visual input 3s before/after saccades to the left for seven of the quantified visual parameters (n=106, 52532 saccades). Bottom row: same as middle row, but for saccades to the right. **d)** saccade trigger computations based on asymmetry (left-right frontal visual field) or signal strength (left+right frontal visual field) of the visual input for the parameters listed in c) (SD out of range). Top row: asymmetry-based computation. Line plots of the mean frontal left visual field (0° to 90°) minus the mean frontal right visual field (0° to –90°) over time for left saccades (blue), right saccades (red), and random frames (grey). Bottom row: signal-strength-based computation. Line plots for adding the mean frontal left visual field (0° to 90°) to the mean frontal right visual field (0° to –90°) over time for left saccades (blue), right saccades (red), and random frames (grey). **e)** Deconstructing the naturalistic visual conditions. The visual features of the virtual landscape were modified, and additional flies were tested. Additional conditions (columns from left to right): The reflective, animated water was replaced with a plain blue texture; the sun was removed; we inverted the intensity while keeping the hue (color information) the same; the height information was removed; and textures were modified to provide a shadeless environment. Top row: panoramic camera renderings for the six visual conditions. Middle row: 2D trajectories of all tested flies (n indicated on top of columns). Scale bars = 20m. Bottom row: heat maps showing density distributions of middle row panels. The color scale was cropped at the 95th percentile of the density data. **f)** Vase plots showing exploration parameters ISI, saccade amplitude, saccade directionality (handedness), and the explored area for each fly of the tested conditions. The data was analyzed using the Wilcoxon rank sum test with asterisks indicating p-values: *p<0.05, **p<0.01, ***p<0.001. **g)** training regression models for determining the hierarchy and influence of visual parameters on saccade triggering, amplitude, and direction. Top row: Models trained on absolute visual asymmetry parameters (left – right frontal visual field). Bottom row: models trained on visual signal strength parameters (left + right frontal visual field). Models with the highest accuracy and lowest RMSE are indicated in color. Bars indicate the coefficient estimate for each visual parameter, with larger bars indicating a stronger influence on the respective saccade parameter. Logistic regression was used for saccade trigger probability (0=no saccade, 1=saccade) and direction (0=left turn, 1=right turn), and linear regression was used for computing the saccade amplitude model. Error bars represent the mean standard error over the 10 cross-validation folds. Asterisks indicate p-values: *p<0.05, **p<0.01, ***p<0.001 (mean over 10 cross-validation folds. **h)** Mean influence of the retinal position of visual input on the saccade parameters. Three linear regression models were trained on the azimuthal visual input (downsampled into 21 bins) up to 500ms before saccades. Coefficient estimates for the respective retinal position were mapped to circular wedges and color-coded. The color scale was cropped at the 95th percentile for better visibility.

### Visual input parameter computations

To compute the panoramic visual input flies experienced, we first re-rendered the panoramic camera output, centered at the fly’s midline, and linked the fly’s rotational movements to the camera’s azimuthal rotation (see Supplemental Video 10, upper panels). From these image sequences, we extracted the mean azimuthal visual input for 9 visual parameters, excluding the blue-sky texture (see Supplemental Video 10, lower panels). Brightness (intensity) input was calculated based on desaturated frames. The blue channel was binarized based on hue information to label only water, and the green channel was binarized based on intensity and hue (dark + green) to include only vegetation. We also extracted the extent of visual motion (optic flow vectors computed using Lucas-Kanade derivative of Gaussian method in MATLAB via opticalFlowLKDoG function, buffered over 3 frames, using the green image channel extracted from the RGB frames to match the chromatic sensitivity of the flies’ motion vision pathway) and it’s mean horizontal and vertical vector components Vx and Vy (Extended Data Fig. 2a,b). Vx and Vy were used to calculate frontal horizontal and vertical visual expansion (when summation of vector components was consistent with outward motion) separately for the left and right visual field as described by: (Tammero & Dickinson, 2002) (see Extended Data Fig. 2a,b). Azimuthal elevation and the sun were also binarized based on intensity and hue information. Furthermore, the mean azimuthal image contrast was defined as the standard deviation of pixel values for each image column.

### Training of regression models for predicting saccade parameters

To compute the influence and hierarchy of visual inputs on saccadic exploration, we trained multivariable logistic and linear regression models on the mean visual input experienced by the flying flies (over land only) up to 500ms preceding saccades for nine visual parameters. The horizontal and vertical components of motion (Vx and Vy) were excluded from the model due to strong correlation with the general motion parameter. We trained separate models for visual processing based on either asymmetry (the mean left frontal visual field minus the right frontal visual field) or signal strength (the sum of visual information across the whole frontal visual field), reflecting two possible strategies for visual signal processing during exploration. We included all saccades from 30 randomly selected flies flying under natural-scene–derived conditions, and all flies flying under modified visual conditions (see Fig. 2e, 61384 saccades total) to expand the range of visual parameters used in training for better model generalization. We computed models for saccade probability (logistic regression, 0 or 1, trained on dataset of 50%/50% saccade/no-saccade frames), saccade amplitude (linear regression, 0° to 180°) and saccade direction (logistic regression, 0 or 1 for left/right) using the mean visual asymmetry or signal strength preceding each saccade as predictor variables and the respective saccade parameter values as behavioral response variables. For logistic regression model training, we used MATLAB’s fitglm function. For linear regression models, MATLAB’s fitlm function was used. Before training, the nine visual parameters were standardized. Training was done using 10-fold cross-validation (CV) to improve generalization. To map the mean influence of the retinal visual input position on the saccade parameters, we trained three regression models (saccade trigger probability, amplitude, and directionality as described above) on the mean whole azimuthal visual input over the last 500ms before saccades were performed (downsampled into 21 azimuthal bins and standardized). Coefficient estimates for the respective retinal positions were ridge-regularized with a fixed L2 penalty, lambda = 10/21.

### Simulating exploration behavior

To test the predictive power of the computed multivariable regression models, we simulated exploration behavior of individual fictive flies in batches of increasing behavioral complexity (see Fig. 4c,d). The simulation framework is explained in detail in Extended Data Fig. 7a. Image features from the panoramic output from the Blender game engine were computed in MATLAB, standardized, and sent to the saccade regression models to predict whether to trigger a saccade, the saccade amplitude, and the saccade direction. First, we simulated a population of blind flies performing random flight with a constant ISI/amplitude based on the median ISI/amplitude of the tested flies (see Fig. 4c,d, Supplemental Video 3). Next, we added the saccade amplitude and directionality models to the simulation, computing visual input deterministically. Then we added a stochastic component to these two models by mapping each model’s predictive output to a linear probability function. E.g., for predicting saccade directionality, if the model output is 0.5 (undecided), amplitude gets chosen randomly. If the prediction is 1, the saccade will be a right turn with 100% probability. If the prediction is 0, the saccade will be a right turn with 100% probability. Values in between were mapped linearly. To predict saccade amplitude, we mapped the directionality model’s output (absolute difference from 0.5) to 0 °– 180 °. This produced more realistic and variable results since the predictions of the saccade amplitude model were mostly close to 40° with only little variation based on visual input. Next, we added the trigger probability model to the previous simulation framework. A saccade was triggered when the model prediction (between 0 and 1) based on visual input was larger than 0.5. We then added circling behavior to the simulation. Flies show an elevated probability to turn in the same direction as the previous saccade (see Extended Data Fig. 7b). We included this memory-based component of exploration behavior by applying the computed mean same-direction probability to the predicted saccade direction based on the time since the last saccade. Lastly, we added individuality to the visual signal processing in the simulation by randomizing the individual saccade trigger thresholds around the mean of 0.5 with a standard deviation of 0.01 (maximum ± 0.3). For simulating behavioral responses within other established visual assays, we adjusted the mean individual saccade trigger threshold: bar tracking assay 0.22, free flight assay 0.5, world map 0.42.

## Results

To investigate how flies respond to simulated naturalistic visual environments, we placed wild-type flies in a tethered flight simulator equipped with a panoramic LED matrix, following a previously established protocol (Mathejczyk et al., 2024) (Fig. 1a). The naturalistic landscape was rendered in real-time using Blender (Blender Foundation, Version 2.79b) in a closed loop guided by a fly’s heading (Supplemental Video 1). As the primary stimulus, we simulated the topography of the Greek island Donousa, using Microsoft Bing satellite imagery in combination with NASA Shuttle Radar Topography Mission (SRTM) elevation data (Fig. 1b; see *Materials and Methods* for details).

### Exploration of Simulated Naturalistic Environments

To assess how flies explore the simulated naturalistic 3D environment, we observed a population of flies (n = 53). Flies quickly explored the whole island and set stable headings by reducing angular velocity when flying over water (Fig. 1c, d, Extended Data Fig. 1b, Supplemental Videos 2 and 3). A preference analysis based on sampling terrain information directly below each fly revealed that flies interacted with the environment in a manner consistent with ecologically relevant terrain selection (Fig. 1e-h). Specifically, the flies strongly preferred flying over land rather than open water (Fig. 1e, Supplemental Video 4). Flies explored the virtual environment quicker and more complete than by random flight (Fig. 1f, Supplemental Video 5). The flies showed a pronounced attraction toward elevated terrain, favoring flight paths toward and over hills (Fig. 1g, Supplemental Video 6). This pattern is consistent with hilltopping, a behavior previously documented in multiple insect taxa, including Diptera and Lepidoptera (Catts, 1979; Pepi et al., 2022; Scott, 1968; Shields, 1967), which increases the likelihood of encountering mates in nature, where high elevations serve as encounter sites. Additionally, the flies preferentially flew over vegetated regions, characterized by a dark, green visual appearance (Supplemental Videos 8 and 9), rather than non-vegetated areas (Fig. 1h, Extended Data Fig. 3a, Supplemental Video 7). The combination of these visual priors allows flies to explore naturalistic environments in a manner consistent with ecologically relevant terrain selection, thereby increasing fitness by increasing the probability of encountering food and mates in natural landscapes.

### Investigating the Cue-Hierarchy Guiding Visual Decision-Making in Saccadic Flight

*Drosophila* flight is characterized by straight trajectories (inter-saccade intervals, ISI) interspersed with rapid heading changes known as ‘body’ saccades (Collett & Land, 1975; Egelhaaf & Kern, 2002; Land, 1999; Land & Collett, 1974) that can either occur spontaneously or be triggered by visual stimuli (Mongeau & Frye, 2017; Tammero & Dickinson, 2002; van Breugel & Dickinson, 2012). Although optic flow and visual expansion have been described as the main visual parameters influencing saccade probability, it remains unclear how flies integrate multiple naturalistic visual stimuli to elicit saccades.

After confirming that flies responded to naturalistic landscapes in a manner consistent with ecologically relevant terrain selection, we sought to quantify saccadic events as important choice points to determine the influence and hierarchy of visual input parameters on saccadic decision-making and, therefore, on small– and large-scale exploration behavior. When analyzing 2D flight trajectories over the virtual island, we detected clear evidence of saccadic exploration, which we quantified using angular-velocity peaks similar to the method described by Mongeau (Mongeau & Frye, 2017). We describe three main saccade parameters influencing exploration behavior: *1. Inter-saccade interval (ISI)* – the time between saccadic events. *2. Saccade amplitude* – the angular magnitude of the turn. *3. Saccade directionality* – whether a fly turns to the left or right (Fig. 2b).

To quantify the visual input perceived before saccades, we re-rendered the panoramic image sequences from the flies’ perspective and extracted the mean azimuthal signal for 11 visual parameters. These parameters included: brightness, green color proportion (binarized vegetation), blue color proportion (binarized water), overall motion (optic flow), the horizon (representing perceived azimuthal elevation distribution), the position of the sun, and image contrast (Supplemental Video 10).

Our analysis included all detected saccades (n=52532), representing 24972 left saccadic turns and 27560 right saccadic turns, and the same number of random samples from non-saccadic episodes. Based on these distinct events, we analyzed how different visual components influence saccade occurrence and directionality (for saccade amplitude dependency, see Extended Data Fig. 3b). This analysis yielded the following results: *Brightness*: Flies turned away from bright regions (based on the left vs. right frontal quadrants of the visual field, L-R), and saccades occurred more frequently in lower-intensity areas (L+R, Fig. 2c1). *Green color*: Flies exhibited a preference for flying toward regions with more green color in the visual field, and saccades were more frequent in green-rich environments (Fig. 2c2). *Blue color*: Flies avoided regions with a higher proportion of blue color in the lower visual field (Fig. 2c3), suggesting a potential aversion to water-like surfaces. Motion components (*total motion, Vx, Vy, horizontal and vertical expansion*): Flies showed avoidance behavior when encountering areas with increased motion, showing that excessive optic flow in the visual field triggered avoidance responses (Fig. 2c4 and Extended Data Fig. 2a, b). Flies displayed an elevated saccade probability after experiencing high levels of frontal horizontal and vertical expansion, but contrary to previous findings (Tammero & Dickinson, 2002), preferred to turn towards visual expansion, probably due to differences in the total amount of optic flow and expansion experienced between the two assays. *Horizon* (elevation perception): Flies were attracted to elevated regions, preferring to turn toward areas where the horizon indicated higher terrain (L-R), and saccades occurred more frequently over elevated terrain (L+R, Fig. 2c5). Position of the *sun*: Using this analysis, no clear L ± R effect on saccadic decision-making was observed (Fig. 2c6). We assume this is due to the sun being represented in a circular compass system within the central complex (Giraldo et al., 2018; Warren et al., 2018), which requires different analytical approaches. *Contrast*: Flies turned away from areas with higher visual contrast (L-R) but performed more saccades in regions with high contrast (L+R, Fig. 2c7).

After analyzing the different image components in the fly’s visual field, we wanted to know whether all image parameters contribute equally to shaping saccadic decisions or whether some components have a stronger influence than others. To de-couple visual parameters and expand the range of visual input for quantifying the hierarchy between the different visual components on saccadic exploration, we generated modified versions of the original landscape (Fig. 2e). Replacing the animated and reflective water with a plain blue surface had no measurable effect on the preference flying over land, confirming that the blue color cue was sufficient to drive water aversion (Fig. 2e2). In the absence of the sun, flies explored significantly smaller areas than when a sun cue was present, consistent with impaired heading stabilization during ISIs (Fig. 2e3, Extended Data Fig. 3c4). Reversing the image brightness while keeping the color information significantly altered directional preferences in relation to the dark green vegetation, indicating that brightness is more important for directional decision making, compared to green color information (Fig. 2e4, see Extended Data Fig. 2c). On a flattened island, flies did not cluster around the previously elevated areas, indicating that elevation specific information is crucial for hilltopping behavior rather than other possibly hill-specific visual properties like intensity or color distributions. Overall, the flattened island became less interesting to the flies, and they started to fly more easily over water. (Fig. 2e5). The shadeless visual scene presents the entire scene in uniform green, drastically reducing contrast and motion cues, disrupting discrimination between land and water. This increased the ISI, expanded the exploration area, and eliminated water avoidance (Fig. 2e6). Furthermore, in the shadeless environment, flies showed a dramatically reduced heading stability during ISIs compared to the naturalistic scene (Extended Data Fig. 3c4).

To quantitatively assess the hierarchy of visual components influencing saccadic decisions, we trained saccade parameter models for ISI, amplitude, and directionality using multivariable regression. We proposed two possible computations within the fly brain for shaping visually guided saccadic parameters: either by computing the asymmetry of the frontal left vs frontal right visual field (L-R) or by summing the signal strength across the whole azimuthal input (L+R), since both computations seemed feasible for triggering visual saccades (see Fig. 2d)

We trained models based on the mean azimuthal visual input up to 500ms before saccades occurred and quantified their predictive power on a held-out dataset containing 10% of saccades. This was repeated ten times (10-fold cross-validation) to increase the robustness of coefficient estimates. The ISI model for predicting the probability of saccade occurrence had slightly better accuracy when based on signal-strength processing (L+R). The probability of saccade occurrence was significantly positively affected by *Green, Blue, Motion*, and *Contrast,* and negatively affected by *Intensity, Horizontal expansion,* and a visible *Sun* (Fig. 2g, left). The saccade amplitude models showed a slightly lower root mean squared error (RMSE) when based on asymmetry processing (L-R). The saccade amplitude is significantly increased by high levels of *Motion* and *Horizontal Expansion* and decreased by high levels of *Brightness, Green, Vertical Expansion,* elevated *Horizon,* and when the *Sun* is visible (Fig. 2g, middle). As expected, the saccade directionality model showed much higher accuracy when based on asymmetry processing (L-R). Flies turn toward elevated levels of *Horizontal* and *Vertical Expansion* and *Horizon*, and turn away from elevated levels of *Brightness, Blue, Motion, Sun,* and *Contrast* (Fig. 2g, right). In accordance with previously described saccade literature, the perceived motion or optic flow is the strongest visual component for shaping the occurrence, amplitude, and direction of visually triggered saccades. However, our analysis reveals a hierarchical structure in which a plethora of visual parameters significantly shape saccade properties. Training models with more sophisticated machine learning techniques also yielded only the predictive power comparable to that of the multivariable regression models used in this study, albeit with less interpretable results (Extended Data Fig. 4).

Next, we trained logistic and linear multivariable regression models on the whole azimuthal visual input (mean of last 500ms before saccades), downsampled into 21 bins, to map the influence of the retinal position of visual input on the three saccade parameters (Fig. 2h, Extended Data Fig. 5). Rather narrow retinal zones in the frontal left and right visual field show the highest influence to visually trigger saccades. The amplitude of saccades is shaped via broader frontal visual input fields. Saccade directionality decisions are influenced via broad lateral visual input across the retina. The most frontal part of the retina exhibits only a weak influence on the three saccade parameters.

### Idiosyncratic motion asymmetry sensitivity thresholds result in individual exploration-exploitation phenotypes

After successfully analyzing and modeling the visual decision-making processes of fly populations, we aimed to determine the extent to which individual differences influence visual exploration strategies. Over a two-day experiment, during which flies were repeatedly tested exploring the virtual island, we observed substantial inter-individuality in flight paths that were consistent over time (Fig. 3a). These differences in flight trajectories were also reflected in two-day correlations of the explored area. Specifically, the total land area explored by each fly across both days correlated with R = 0.39.

**Fig. 3:**
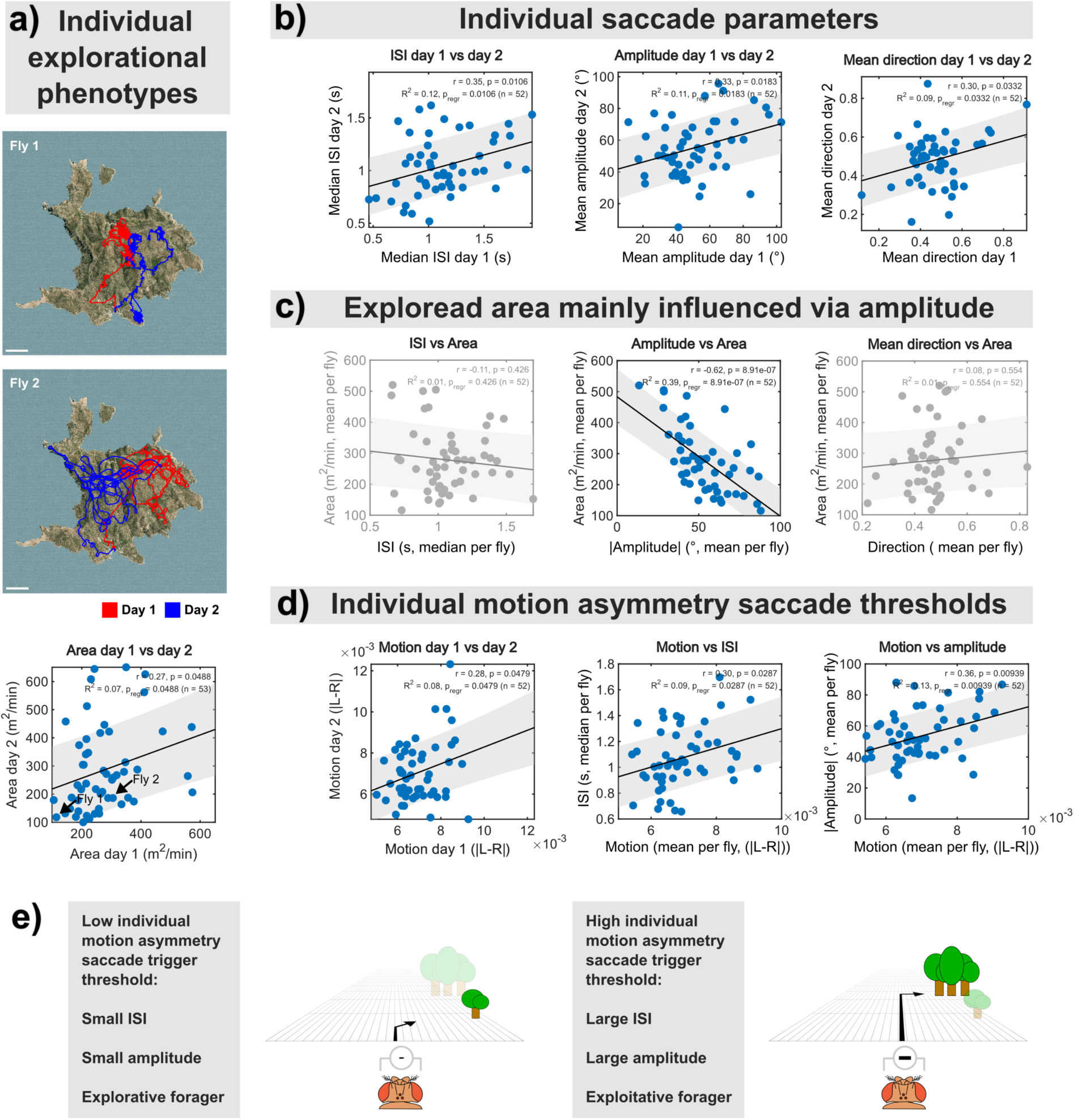
Exploration VS. exploitation – Individual differences in visual signal processing affect individual explorational phenotypes. **a)** Exploration strategy as an individually stable trait. Top two panels: two-day trajectories from two exemplary flies (day 1: red, day 2: blue) illustrating different exploratory phenotypes, with fly 1 showing a rather explicatory search strategy and fly 2 being more explorative (larger explored area) on consecutive days. Scale bar: 20m. Bottom: Scatter plot of the area the individual flies explored on two consecutive days. The data from fly 1 and fly 2 from the upper panels are marked with black arrows. n=53. The grey shaded area marks the 95% CI. **b)** Individual saccade parameters are individually stable traits. Scatter plots of individual day 1 vs day 2 saccade parameters. Left: Median ISI. Middle: mean saccade amplitude. Right: mean saccade direction (0=left, 1=right). Filter threshold >200 saccades per trial. n=52. Grey-shaded areas mark 95% CIs. **c)** The explored area is mainly influenced by saccade amplitude. Non-significant panels are shaded grey. Left: Median ISI vs area. Middle: mean saccade amplitude vs area. Right: mean saccade direction (0=left, 1=right) vs area. Filter threshold >200 saccades per trial. n=52. **d)** Flies show individual motion asymmetry saccade thresholds, which influence individual ISI and saccade amplitude. Left: the absolute mean asymmetry (left minus right frontal visual field) of motion (optic flow) perceived up to 500ms before saccades is plotted for day 1 vs day 2 trials. Some flies already perform saccades after seeing low visual motion asymmetry, while others only after seeing high visual motion asymmetry. Middle and right: individual motion asymmetry saccade thresholds influence individual ISI and saccade amplitude. Filter threshold >200 saccades per trial. n=52. **e)** Graphical summary of Fig. 3. Left: a fly with a low motion asymmetry saccade trigger threshold is more likely to perform saccades already at rather low asymmetry levels in the visual field, leading to shorter inter-saccade intervals on average. However, such a fly would also show larger saccade amplitudes, leading to a smaller but more thoroughly explored area of land (exploitative phenotype). Right: a fly with a high visual motion asymmetry saccade trigger threshold is more likely to perform saccades only at rather high asymmetry levels, leading to longer inter-saccade intervals on average. Such a fly would also perform smaller saccade amplitudes, leading to a larger but less thoroughly explored area of land (explorative phenotype). **a-d)** Correlations were quantified using Pearson’s correlation coefficient (r).

Also, several saccade properties are inter-individually variable and consistent over time. The individual ISI over land showed a correlation of r = 0.35 (Fig. 3b, left), while saccade amplitude showed a correlation of r = 0.33 (Fig. 3b, middle). Similarly, mean individual saccade directionality (handedness) correlated with r = 0.3 (Fig. 3b, right). Furthermore, we found that the explored area is mainly influenced by the saccade amplitude, with a strongly negative correlation (r = –0.62). This means flies performing larger amplitudes tend to explore smaller areas very thoroughly, while flies performing small amplitude saccades explore larger areas, but less thoroughly (Fig. 3c). On a population level, this ensures a balance of full island exploration and thorough exploitation of resources (vegetation). Since saccades can be visually triggered, we next asked whether we could identify individual, persistent visual saccade-trigger sensitivities in the dataset. We analyzed the mean visual input based on asymmetry (L-R) and signal strength (L+R) at which flies performed saccades for the eight main visual parameters (excluding sun) and only found significant individual correlation with the symmetry processing of motion (r = 0.28, Fig. 3d left, Extended Data Fig. 6a, b). Flies that performed saccades already at low visual motion asymmetry levels on day 1 tend to do so also on day 2, whereas flies performing saccades only after perceiving high visual motion asymmetry on day 1 also display these individual motion asymmetry sensitivity thresholds on day 2. Furthermore, these individual motion asymmetry thresholds also correlate with the individual median ISI and mean saccade amplitude during exploration behavior (Fig. 3d middle, right). These findings demonstrate that idiosyncratic motion asymmetry sensitivity thresholds predict explorational phenotypes (explorative vs exploitative) mainly via affecting the individual mean saccade amplitude. Flies with lower motion asymmetry thresholds (i.e., higher sensitivity to motion asymmetry) exhibited shorter ISIs and smaller amplitudes, leading to broader exploration patterns. Conversely, flies with higher motion thresholds (i.e., lower motion sensitivity) exhibited longer ISIs and larger saccade amplitudes, resulting in narrower, more exploitative exploration patterns (Fig. 3e).

### An exploration strategy that maximizes resource and mate encounters

Lévy walks, or Lévy flights, are stochastic movement processes with heavy-tailed (typically power-law) step-length distributions, producing occasional long displacements among many short movements. This movement pattern leads to super-diffusive behavior, enabling faster spread over time than random Brownian motion. This pattern has been observed across various natural systems, including animal foraging behavior. The Lévy flight foraging hypothesis posits that Lévy walks have evolved to optimize search efficiency when resources are sparsely distributed, a claim supported by empirical evidence across a wide range of species (Campeau et al., 2022).

We analyzed the Lévy properties of the ISI distribution of flies flying over the naturalistic Island. Lévy flight is characterized by a long tail in the step-length (ISI) distribution following a power law with an exponent α between 1 and 3, with an optimum at α = 2. Flies exhibited exploration behavior with Lévy properties close to the mathematical optimum (Fig. 4a, left). An analysis of individual Lévy properties revealed that most flies performed Lévy-like search behavior on the island, with some flies being more explorative (α close to 1) and others being more exploitative (α close to 3), and a population median close to the mathematical optimum of α = 2 (Fig. 4a, right). Since flies must not only find food sources but also mates in nature, we further investigated whether hilltopping behavior would increase the likelihood of encountering potential mating partners by reducing inter-fly distances within the fly population. Due to the relatively low spatial resolution of the *Drosophila* visual system and the small size of the flies, finding mates by vision, especially in a complex natural scene, is challenging. However, we find that hilltopping behavior reduces inter-fly distances, drastically increasing the chances of encountering mates (Fig. 4b).

**Fig. 4:**
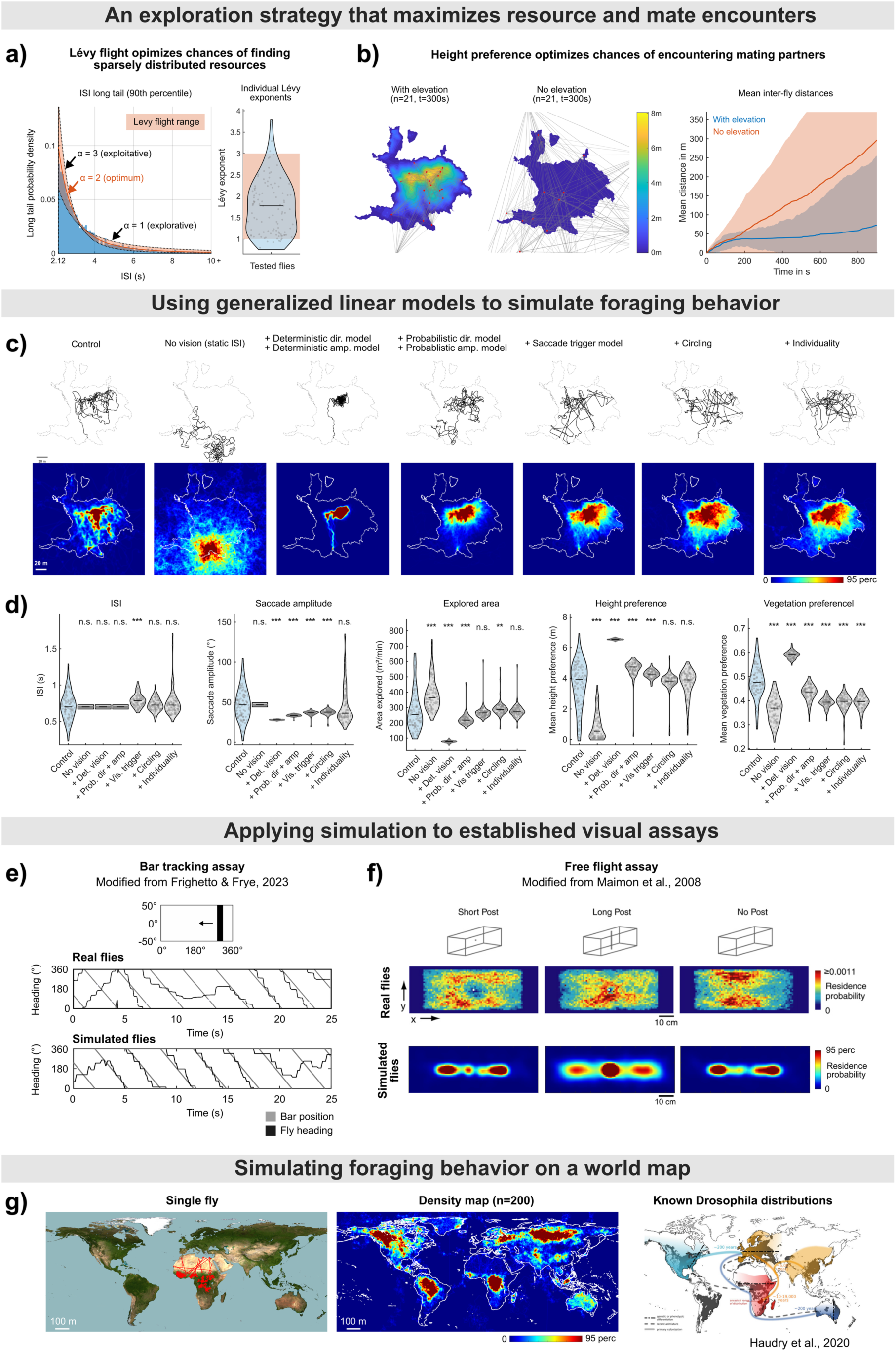
Optimized exploration strategy and simulation of exploration behavior. **a)** Flies perform optimal Lévy flight, optimizing chances of finding sparsely distributed food sources. Left: Histogram of the long tail (90th percentile, 4403 saccades, xmin=2.12s) of the ISI distribution when flying on land under naturalistic visual conditions. Lévy flight is characterized by the ISI long tail distributions following a power-law with a power-law exponent α between 1 (more long inter-saccade intervals) and 3 (less long inter-saccade intervals), with an optimal search efficiency around α=2. The respective power-law functions were fitted to the long tail of the ISI distribution and plotted alongside the ISI tail histogram. Right: Vase plot showing individual power-law exponents computed using the Maximum Likelihood Estimation analysis software described in (Humphries et al., 2010). Lévy range of power-law exponents α is shaded in red. **b)** Visually guided height preference optimizes chances of encountering mating partners through minimizing inter-fly distances during exploration. Left: fly population distribution 300s after being released in a naturalistic environment containing elevation data. Grey lines link each fly to all other flies of the population. n=21 (randomly selected from the fly population described in Fig. 1). Middle: same as left but for flies flying on a flat island without elevation data (see Fig. 2e, n=21). Right: Mean inter-fly distance over time for the fly populations exploring the island shown in the first two panels. Shaded areas indicate ±1 SD**. c)** Applying generalized linear models (see Fig. 2g) to simulate and dissect exploration behavior and its underlying components. Top row: 2D trajectories of single flies of each test condition (left: real fly control data, to the right: simulated flies). Scale bar: 20m. Bottom row: heat maps showing density distributions of top row panels (n=106 flies per group). The color scale was cropped at the 95th percentile of the density data. Scale bar: 20m. **d)** Vase plots for five key exploratory behavioral parameters for control and simulated flies, n=106 per group. The data was analyzed using the Wilcoxon rank sum test with asterisks indicating p-values: *p<0.05, **p<0.01, ***p<0.001. **e-f)** Applying generalized linear saccade models to established visual assays/stimuli. **e)** Bar tracking assay. Top: Videos of a black vertical bar (width 30°) moving counterclockwise with 112.5°/s (modeled after stimulus described in (Frighetto & Frye, 2023)) were used as visual stimuli within the simulation framework (see Extended Data Figure 7a) instead of the panoramic camera renderings. Middle: Heading of a real tested fly (black line) in response to the moving bar positions (grey). Bottom: Like the middle panel, but for a simulated fly. The model produces results very similar to real fly behavior, even though the regression models were not trained on this type of visual stimulus. **f)** Free flight assay. We modeled the free-flight assay described in (Maimon et al., 2008) (see Extended Data Fig. 7c), adding either a short or a long back cylindrical post, or no post, to the arena center (top panel, row). Middle: original flight trajectory heatmaps from (Maimon et al., 2008), showing short post avoidance and long post attraction in real flies. Bottom: Heatmaps of simulated fly trajectories showing clustering around a long post but no avoidance of the short post, since models were not trained on the vertical extent of visual input. n=120 flies per panel (15min, 80Hz sampling rate). **g)** Simulating exploration behavior on a world map (scaled down to 1km x 2km). Left: Trajectory of a single representative fly showing an increased saccade frequency over vegetated regions and only few saccades over desert regions. Middle: Heatmaps of the 2D trajectories of 220 simulated flies. The color scale was cropped at the 95th percentile of the density data for better visibility. Right: Map of known *Drosophila* distributions from (Haudry et al., 2020), showing similar fly distributions to the simulation in the middle panel (e.g. clustering in central Africa and low detection probability in the Sahara Desert).

### Model-based simulation replicates flight statistics

To validate the applicability of the trained saccade parameter models, we simulated *in silico* flies exploring the landscape using these models and compared their behavioral statistics to those of the real fly populations (Fig. 4c, d).

We began with the simplest simulated flies, which had a constant ISI based on *in vivo* fly data and made directional decisions at random, independent of visual input. These simulated flies dispersed randomly, resulting in the highest flight path density near the starting point, with decreasing density farther away (Fig. 4c2). In the next iteration, we added saccade amplitude and directionality prediction to the simulation by mapping the output of these respective models deterministically to saccadic behavior based on visual input (Fig. 4c3). These simulated flies did not explore the entire island; instead, most flew directly to the area that elicited the strongest visual attraction, effectively functioning as a visual trap. In the virtual representation of Donousa, this was the hill region in the center of the island. Next, we simulated flies using a probabilistic vision model (Fig. 4c4). This led to flies exploring the island more completely. Next, we added the saccade trigger model to the simulation (Fig. 4c5). These simulated flies now triggered saccades in response to visual input and explored an even larger area than in the previous version. To further refine the simulation, we added circling behavior based on the time since the last saccade (Fig. 4c6, Extended Data Fig. 7b). An observation from our behavioral data was that flies exhibit a directional bias, where a rightward saccade increases the likelihood of another rightward saccade (same for left saccades), resulting in characteristic circling behavior. When we incorporated this saccadic memory, flies exhibited the same height preference as the real flies in the experiment. Lastly, we incorporated individuality into the simulation by adding noise to the individual saccade trigger threshold (Fig. 4c7). This increased the overall variance in behavioral parameters by either making simulated flies more exploitative or more exploratory.

After confirming that the simulated flies accurately replicated *in vivo* fly behavior in the naturalistic environment, we tested their performance in novel visual scenarios not included in the training dataset. The first test involved presenting the flies with a dark vertical bar moving azimuthally against a bright background, a classical stimulus used in optomotor response experiments (adapted from (Frighetto & Frye, 2023)). Despite having never encountered such a visual stimulus before, the simulated flies responded almost identically to real flies, following the bar rotation probabilistically. This demonstrated the generality of the saccadic models presented in this study (Fig. 4e). Next, we tested the simulated flies flight responses in a virtual arena containing either a long or a short post or no post as described in (Maimon et al., 2008) (Fig. 4f). The simulated flies exhibited less noisy dispersal and clustered around the long post, similar to *in vivo* flies, but did not show the strong avoidance of the short post.

Given the robustness of our *in silico* model and simulations, we investigated whether visual-driven exploration models could provide insights into real-world fly dispersal and distribution patterns. Using a scaled-down high-resolution map of Earth (https://en.wikipedia.org/wiki/File:Blue_Marble_2002.png), we analyzed the global detection probability of simulated flies and compared these predictions with measured *Drosophila* distributions (Haudry et al., 2020). Surprisingly, the simulated dispersal map closely resembled the empirical distribution of wild *Drosophila* populations (Fig. 4g), e.g., showing strong clustering in central Africa and low densities over the Sahara Desert.

Together, our results demonstrate that *Drosophila’s* saccadic flight behavior is controlled by a hierarchical integration of visual cues, in which simple, interpretable sensory decision rules, modulated by stable individual differences, are sufficient to generate efficient exploration of complex natural landscapes across spatial scales. By linking moment-to-moment visual processing at the level of individual saccades to emergent population-level search efficiency, our framework provides a mechanistic bridge between neural computation, behavioral individuality, and ecological performance. More broadly, our findings suggest that scalable exploration in complex environments can arise from compact, stochastic sensorimotor rules rather than detailed internal representations of the world.

## Discussion

### Short highlight of key points

The results presented in this paper confirm that saccadic exploration can be accurately simulated using data-driven approaches, capturing both population-level trends and individual-level differences. Our work provides a computational framework that may be useful for studying insect movements and unmanned autonomous vehicles under natural conditions. In essence, our findings demonstrate that insect vision-based exploration behavior can be modeled by relatively simple hierarchical and idiosyncratic signal-processing mechanisms, enabling flies to efficiently explore complex natural environments and increase the likelihood of encountering resources and potential mating partners.

### A minimalistic world model guides exploration within naturalistic, large-scale landscapes

This study was built upon the growing body of research investigating complex visually guided behaviors by leveraging a high-resolution, naturalistic 3D virtual reality environment. While many studies of visually guided behaviors have successfully employed low-dimensional visual cues, the experimental framework presented here bridges well-controlled vision-based laboratory studies and real-world ecological observations, allowing us to examine how flies explore simulated large-scale naturalistic landscapes by integrating a complex set of visual parameters.

Our results demonstrate that visually guided large-scale exploration in *Drosophila* is structured, not random, with macroscopic path structure emerging from cue-dependent saccadic decisions. Although the flies had never flown in nature, they showed innate visual priors well-suited to successfully exploring complex naturalistic terrains. Flies preferred elevated, vegetated areas while avoiding flying over water. These simple visual priors are consistent with a minimalistic visual world model hypothesis and are expected to enhance the flies’ fitness in nature by biasing exploration toward ecologically favorable terrain, thereby increasing the likelihood of encountering food sources and mates. In this study, we use the term minimalistic visual world model in a strictly operational sense: a behaviorally efficacious, structure-preserving, and ecologically compressed internal representation (Yildirim & Paul, 2024) of a naturalistic combination of visual scene statistics that is sufficient for guiding adaptive exploration of novel natural landscapes without implying higher cognitive functions like planning, cognitive maps, or semantic scene understanding. We use this term because flies not only showed reflex-like responses to low-dimensional visual cues, but instead exhibited stable, adaptive, and context-dependent saccadic behavior in response to a complex composition of naturalistic visual scene parameters. Consistent with this interpretation, we could quantitatively link specific visual scene statistics to context-dependent exploration behavior and naturalistic terrain selection.

While prior studies successfully connected natural scene statistics to contrast coding in early fly visual interneurons (Laughlin, 1981), our results show that multivariable natural scene structure shapes naturalistic and biologically relevant saccadic responses and stable individual exploration strategies at the behavioral level. Our findings are also consistent with previous studies showing that *Drosophila* exhibits innate and state-dependent spectral preferences, with color processing entangled with brightness and behavioral context (Lazopulo et al., 2019; Schnaitmann et al., 2013). Likewise, the attraction to hills observed in flies exploring the virtual island is consistent with hilltopping (Catts, 1979; Pepi et al., 2022; Scott, 1968; Shields, 1967), a behavior previously not reported in *Drosophila*. Small insects, such as flies, are hard to track in natural environments. Therefore, the assay described in this study provides a means to quantify ecologically relevant behaviors such as hilltopping, terrain selection, and even large-scale migratory movements at the population level.

### Sensory processing and signal hierarchy

By mapping visual scene statistics to saccadic behavioral output, we trained predictive models for saccade probability, amplitude, and direction, revealing a hierarchy of the influence of visual input parameters on saccadic exploration, with motion cues dominating saccadic decisions. A hierarchy of visual cues also emerged from systematically modifying the simulated environment, confirming that certain visual factors exert stronger influences on saccades than others. For instance, reversing brightness while keeping color information the same reversed saccade directionality decisions, indicating that brightness processing is more important for choosing turn direction than color vision. These results mirror previous findings that insects integrate multiple visual cues hierarchically to optimize flight trajectories (Giger & Srinivasan, 1997). At the mechanistic level, this cue hierarchy is also in accordance with prior work showing that saccadic flight maneuvers are strongly gated by perceived motion and visual expansion (Mongeau & Frye, 2017; Reiser & Dickinson, 2013; Tammero & Dickinson, 2002). The flies in our assay were attracted to frontal visual expansion, although Tammero & Dickinson (2002) describe an aversion to frontal visual expansion during flight. We speculate that this is due to differences in optic flow across the two visual assays, gating distinct behavioral responses in a context-dependent manner.

We demonstrated the importance of stochasticity in saccadic decision making, which, in contrast to rigid, reflex-like responses, prevents flies from getting stuck in visual ‘traps’. Furthermore, stochastic opto-motor decision-making increases the chances of evading predatory attacks. In this regard, it was previously shown that fly escape circuits in *Drosophila* can integrate multiple visual looming features and produce stochastic, context-dependent motor output (von Reyn et al., 2017).

### Individual differences in signal processing shape large-scale exploration on a population level

Multi-day testing revealed stable and repeatable differences in asymmetry-based motion processing, shaping individual exploration-exploitation strategies. This key finding aligns well with previous studies demonstrating that consistent behavioral individual differences exist even under controlled genetic backgrounds and environments (Ayroles et al., 2015; Buchanan et al., 2015; Kain et al., 2012; Linneweber et al., 2020).

Mechanistically, asymmetry-based exploration phenotypes are consistent with prior studies showing that fly individuality can emerge from stable inter-individual differences in neural circuit structure and gain control. Work on individually stable left-right choice behavior points to central brain circuits driving locomotor handedness, while other studies show that idiosyncratic neuromodulation and sensory processing through variable neuronal circuits can generate persistent individual response patterns (Buchanan et al., 2015; Honegger et al., 2020; Linneweber et al., 2020).

From an ecological and evolutionary perspective, a mixture of exploration–exploitation phenotypes in the population functions as a bet-hedging strategy, which might be advantageous in variable, rapidly changing natural landscapes where resources are scarce and randomly distributed, since diverse individual strategies within a population may sample resources more robustly. This is in accordance with previous studies showing adaptive bet-hedging in sensory preferences and aligns with theoretical work linking stable behavioral variation to ecological performance and fitness outcomes (Kain et al., 2015; Smith & Blumstein, 2008; Wolf & Weissing, 2012).

We furthermore found individual exploration-exploitation variability at the population level by quantifying Lévy properties of 2D trajectories, indicating that flies don’t just follow a strict exploratory routine, but rather employ a spectrum of search strategies suitable for exploring large-scale territories while efficiently exploiting resources. This is in accordance with prior work demonstrating Lévy-like flight characteristics during search behavior in *Drosophila* (Reynolds & Frye, 2007). Lévy-like foraging has previously been shown to be strongly content-dependent, with prior work even demonstrating behavioral switching between Lévy-like and Brownian movements across different terrains (Humphries et al., 2010; Palyulin et al., 2014). However, the Lévy-like foraging variability observed in this study indicates that flies combine individual– and context-dependent locomotor foraging strategies that balance large-area exploration with local resource exploitation in heterogeneous natural environments.

### Using predictive models for simulating saccadic behavior

The compact, data-driven modeling framework presented in this study successfully reproduced key exploratory and trajectory statistics by predicting ISI, saccade amplitude, and direction from recent visual input. Our finding that saccade probability is better predicted by signal strength (L+R) computations, while asymmetry-based (L-R) modelling performs better for amplitude and directionality predictions, indicates different neural substrates processing timing and turning properties. Signal strength-based (L+R) saccade triggering fits well with prior studies showing that direction-sensitive motion signals are computed in T4/T5 pathways and are mapped to motion-axis-specific lobula plate layers, serving as a robust neural substrate suitable for global motion-strength estimates that could gate visual saccade triggering (Haag et al., 2017; Maisak et al., 2013). In contrast, saccade amplitude and directionality decisions are likely modified via bilateral opponent processing (L-R) and descending motor control pathways. Prior work successfully described descending interneurons that integrate wide-field motion signals and project to bilateral flight motor circuits, driving saccadic turn maneuvers (Namiki et al., 2022; Schnell et al., 2017; Suver et al., 2016).

Furthermore, additional motor control is also likely provided via central-complex heading networks, integrating self-motion and landmark cues and coupling internal heading representations to saccadic turn behavior. This framework offers a mechanistic basis for cue-dependent control of large-scale saccadic exploration (Kim et al., 2017; Seelig & Jayaraman, 2015; Shiozaki et al., 2020; Turner-Evans et al., 2020). Our modelling approach translated well to other visual assays, indicating that the compact, hierarchical, vision-guided saccade framework described here captures core signal processing that generalizes across different visual assays and spatial scales. Surprisingly, the simulations also predicted large-scale fly dispersal on a world map. However, this result should be interpreted cautiously, because our simulation omits many real-world factors, such as wind, learning, and olfaction, and therefore provides a proof of principle rather than a direct reconstruction of natural dispersal. Furthermore, Lévy-like exploration patterns and basic elements of the underlying behavior may be close to scale-invariant, meaning that this special distribution of ISI lengths enables efficient exploration of natural terrain across a wide range of visual scales and spatial frequencies, allowing us to extract general properties even from scaled-down environments. The methods and findings described in this study provide a novel approach not only for computing mechanistic models of visually guided saccadic navigation but also for studying and predicting large-scale insect population dynamics under natural conditions. Furthermore, our methods could be an efficient way to control UAVs in various natural terrains. However, natural cues that influence insect exploration and migration are not limited to vision alone. Among others, wind drift (Leitch et al., 2021), olfactory cues, and temperature gradients can also influence flight trajectories. Extending the stimulus space in future studies may allow a deeper understanding of the hierarchy and computations not only of vision but also of a wide variety of sensory modalities. Future studies will also aim to map specific model parameters to previously described neural circuits using modern *Drosophila* genetic tools and whole-brain connectome data (Dorkenwald et al., 2024).

### Discussion Summary

In summary, our findings provide new insights into the visual basis of insect exploration behavior, demonstrating that *Drosophila* relies on a structured hierarchy of visual cues to make flight decisions. The results presented in this study show how the integration of multiple visual cues, in combination with individual motion sensitivities, leads to efficient exploration of naturalistic landscapes at the population level. These findings bridge the gap between controlled laboratory studies and real-world observations, providing a framework for future research on insect movement ecology. Moreover, our *in-silico* fly simulations suggest that visual-driven models can effectively predict insect dispersal patterns, with potential applications in ecology and conservation science. Future research will investigate the neural mechanisms underlying these behaviors, as well as the potential ecological relevance of individual navigation strategies in natural environments.

## Acknowledgements

The authors thank the Bloomington Stock Center for fly stocks and reagents. This work was supported by the Deutsche Forschungsgemeinschaft (DFG) through the DFG research unit 5289 RobustCircuit (G.A.L) through grants LI 2640/1-1, FOR5289 LI 2640/2-1 (G.A.L.), and with support from the Fachbereich Biologie, Chemie & Pharmazie of the Freie Universität Berlin, as well as the Division of Neurobiology at Freie Universität Berlin. We thank all members of the Linneweber lab, Randolf Menzel, Mathias Wernet, and Robin Hiesinger, for helpful discussions. We also thank Nick Humphries and David Sims for helping with the Lévy analysis.

## Author Contributions

Conceptualization, T.F.M. and G.A.L.; Methodology, T.F.M. and G.A.L.; Investigation, T.F.M.; Resources, T.F.M. and G.A.L.; Supervision, G.A.L.; Funding Acquisition, G.A.L

## Supplemental figure legends

**Extended Data Fig. 1:**
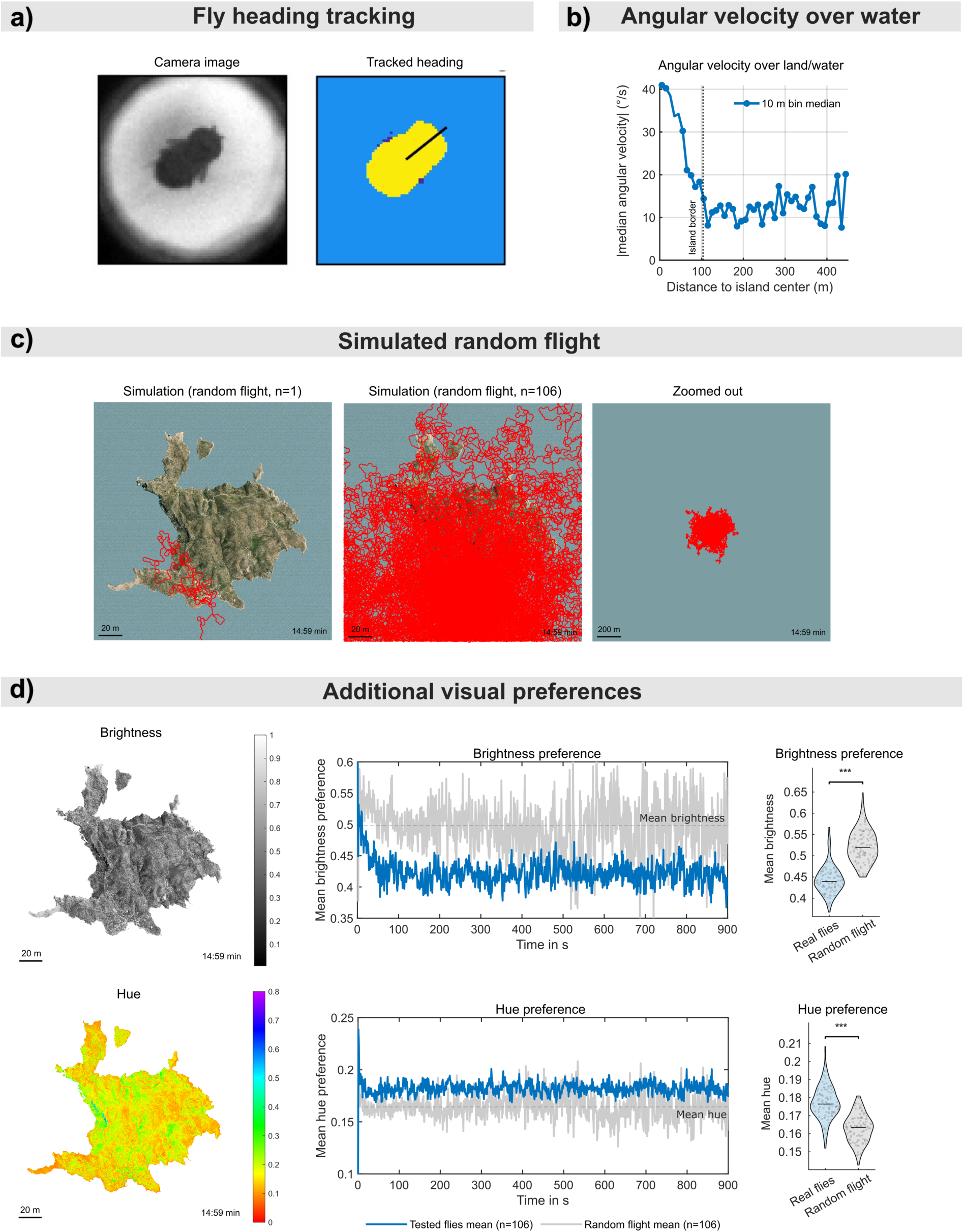
a) Tracking of fly heading. Left: Camera image. Right: The fly’s body is binarized from the background, and its heading is extracted by fitting an ellipse around the fly’s body pixels. Angular ambiguity is solved based on anterior-posterior symmetry (fewer pixels in the head region). **b)** Flies strongly reduce angular velocity over water. Median absolute angular velocity of the tested fly population (n=106) depending on distance to the island center, binned in 10m steps. The 99-percentile island border is marked with a vertical dotted line. **c)** Simulation of random flight. We simulated 106 ‘blind’ flies performing random flight (random directionality, static individual ISI, and amplitude based on the median of measured real-fly distributions). Left: 2D trajectory of a single simulated blind fly. Scale bar: 20m. Middle: 2D trajectories of all simulated blind flies (n=106). Scale bar: 20m Right: Zoomed out map region from the middle panel. Scale bar: 200m. **d)** Visual preferences for brightness and hue (color). Left: Maps of the island showing brightness and hue information of the landscape. Scale bars: 20m. Middle: Mean brightness and hue preferences of tested flies (blue) and simulated flies performing random flight (grey). Mean brightness and hue of the island are marked with dotted lines. Flies prefer to fly over dark and green patches of land (vegetation). Right: Mean individual brightness and hue preferences for real flies (blue) and random flight (grey). The data was analyzed using the Wilcoxon rank sum test with asterisks indicating p-values: *p<0.05, **p<0.01, ***p<0.001.

**Extended Data Fig. 2:**
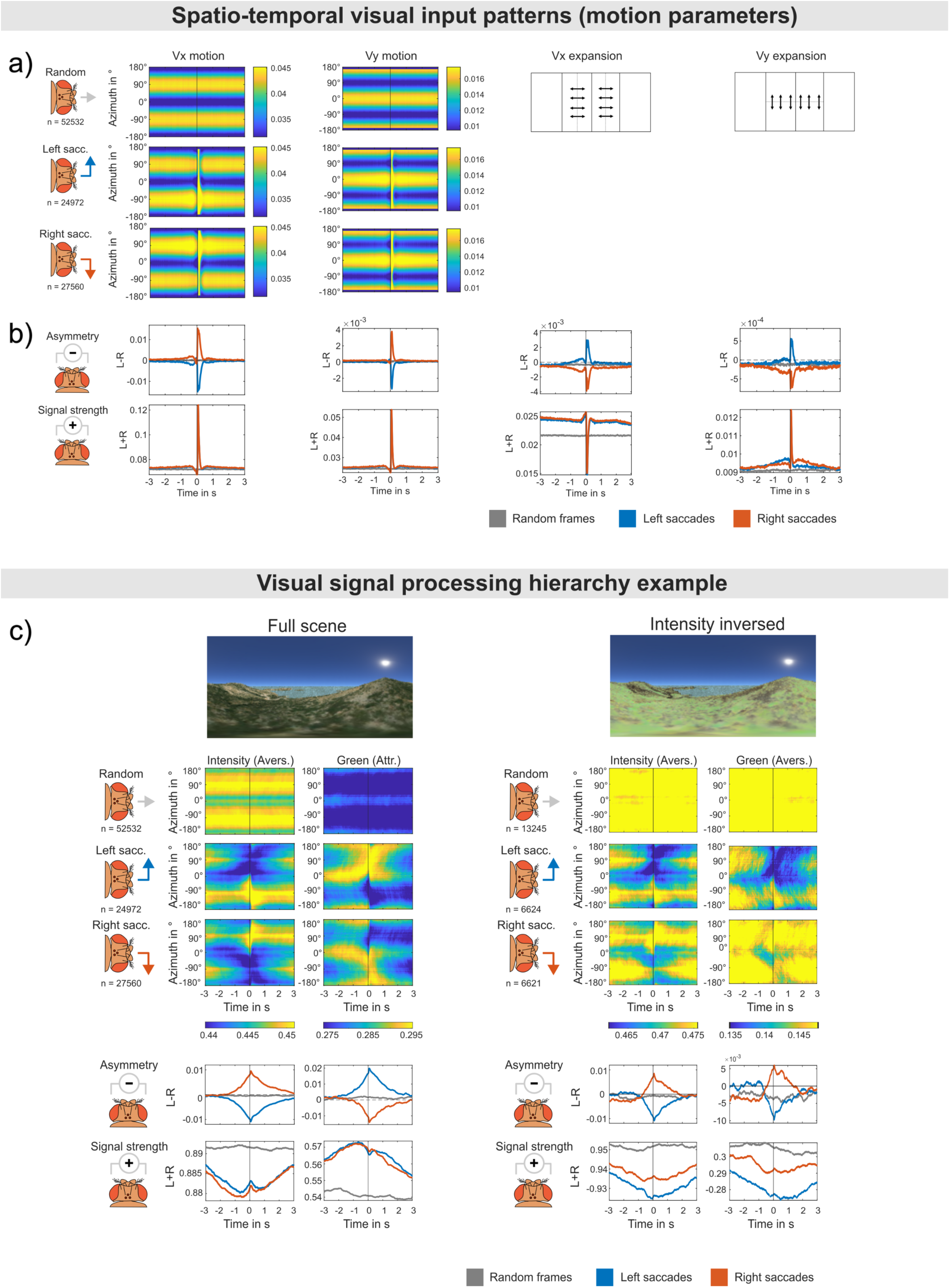
Spatio-temporal motion component patterns and visual hierarchy example. **a)** Averaged spatio-temporal azimuthal visual input patterns of non-saccadic periods and all detected saccades aligned at time point 0s (land-based saccades only) for the horizontal (Vx) and vertical (Vy) components of the optic flow vector field. Schematic vector field processing is indicated for horizontal and vertical frontal expansion based on the visual panorama. For a detailed description of the plotting scheme, see Fig. 2c. **b)** Possible saccade trigger computations based on asymmetry (left-right frontal visual field) or signal strength (left+right frontal visual field) of the visual input for the motion parameters listed in a). For a detailed description of the plotting scheme, see Fig. 2d. **c)** Intensity processing is more important for determining saccade direction than color information. Spatio-temporal patterns and asymmetry/signal strength computations in a naturalistic environment (left, data from Fig. 2c, d, n=106) and in the same environment with inverted intensity, but the same color information (right, n=32). With inverted brightness, flies still turn away from high brightness levels, as in a natural-scene–derived environment, but also turn away from green color information, demonstrating a higher importance of visual brightness in making directional decisions than of color information. For a detailed explanation of the plotting scheme, see Fig. 2c, d).

**Extended Data Fig. 3:**
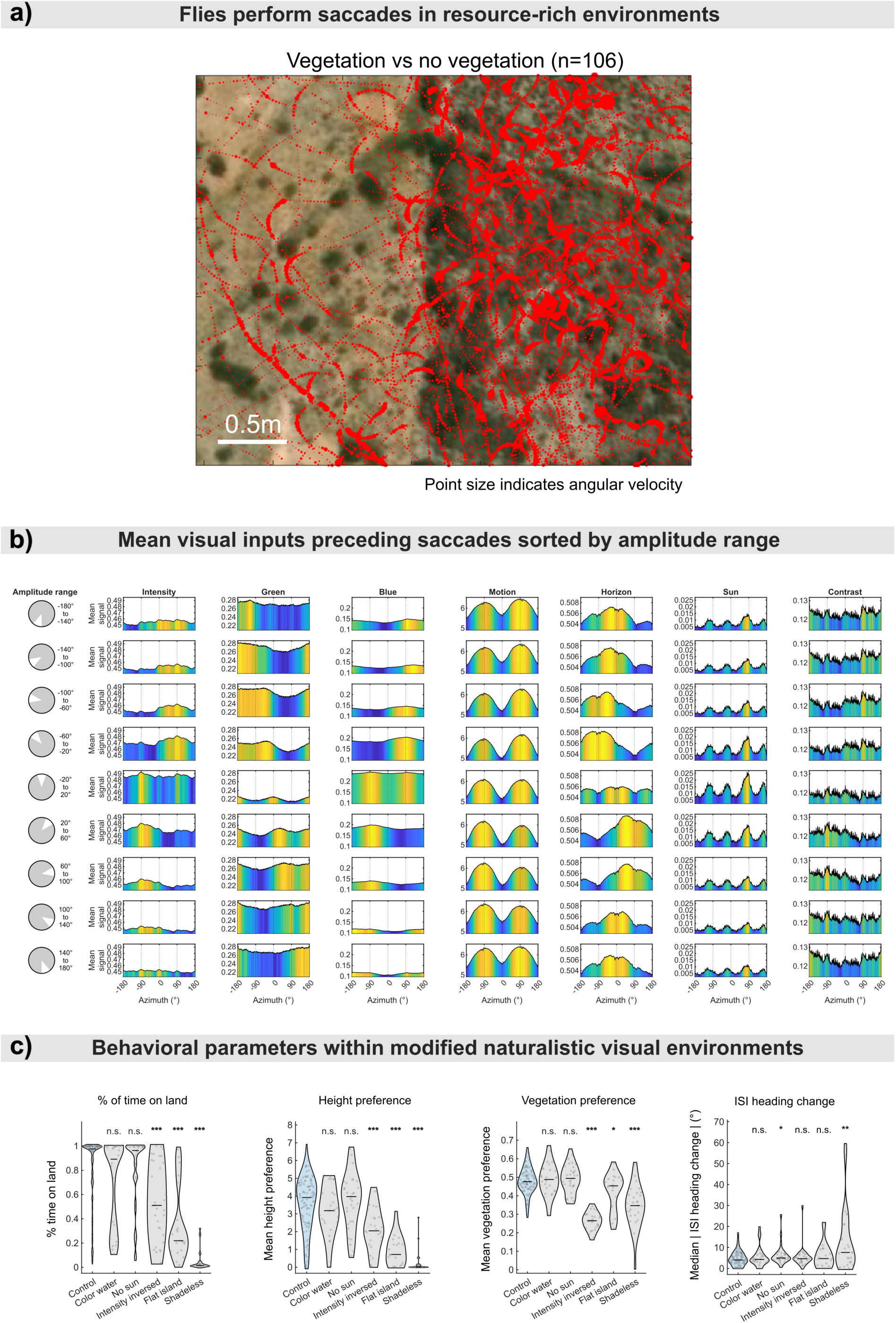
Visually guided saccade properties lead to vegetation exploitation. **a)** Flies perform saccades over resource-rich environments. Zoomed-in section of flight trajectories from Fig. 1c (middle) showing flight responses over high vs low-vegetation areas. Each marker point (red) indicates a fly’s (n=106) flight position. The marker size indicates the absolute measured angular velocity at that position. Flies perform more saccades over vegetation, increasing the time spent flying over vegetated areas and the chances of finding food. Scale bar: 0.5m. **b)** Mean visual inputs preceding saccades sorted by saccade amplitude range. Each row shows the mean visual panorama experienced up to 500ms before saccades were performed (data from Fig. 2c), sorted by saccade amplitude range (–180° to 180°). Each column shows data for the 7 analyzed visual parameters. The color scale range was set to min/max values of each parameter to show possible azimuthal shifts of parameter strength preceding saccades over different ranges of saccade amplitudes. E.g., green visual input (vegetation) invokes targeted saccade amplitudes depending on the azimuthal position of the most vegetation (shift of yellow peaks) across amplitude ranges. On the other hand, azimuthal brightness information appears to be processed within the fly brain, rather than coarse left vs right visual field comparisons (with peaks/valleys around ±90°), regardless of the exact azimuthal position of bright/dark areas. **c)** Overview of additional behavioral parameters for the modified naturalistic environmental stimuli. Vase plots show individual behavioral parameters computed for each fly of the tested conditions. The ISI heading change indicates the individual median absolute angular drift between the start and end of ISIs, averaged over the first and last 10 frames, respectively. The data was analyzed using the Wilcoxon rank sum test with asterisks indicating p-values: *p<0.05, **p<0.01, ***p<0.001.

**Extended Data Fig. 4:**
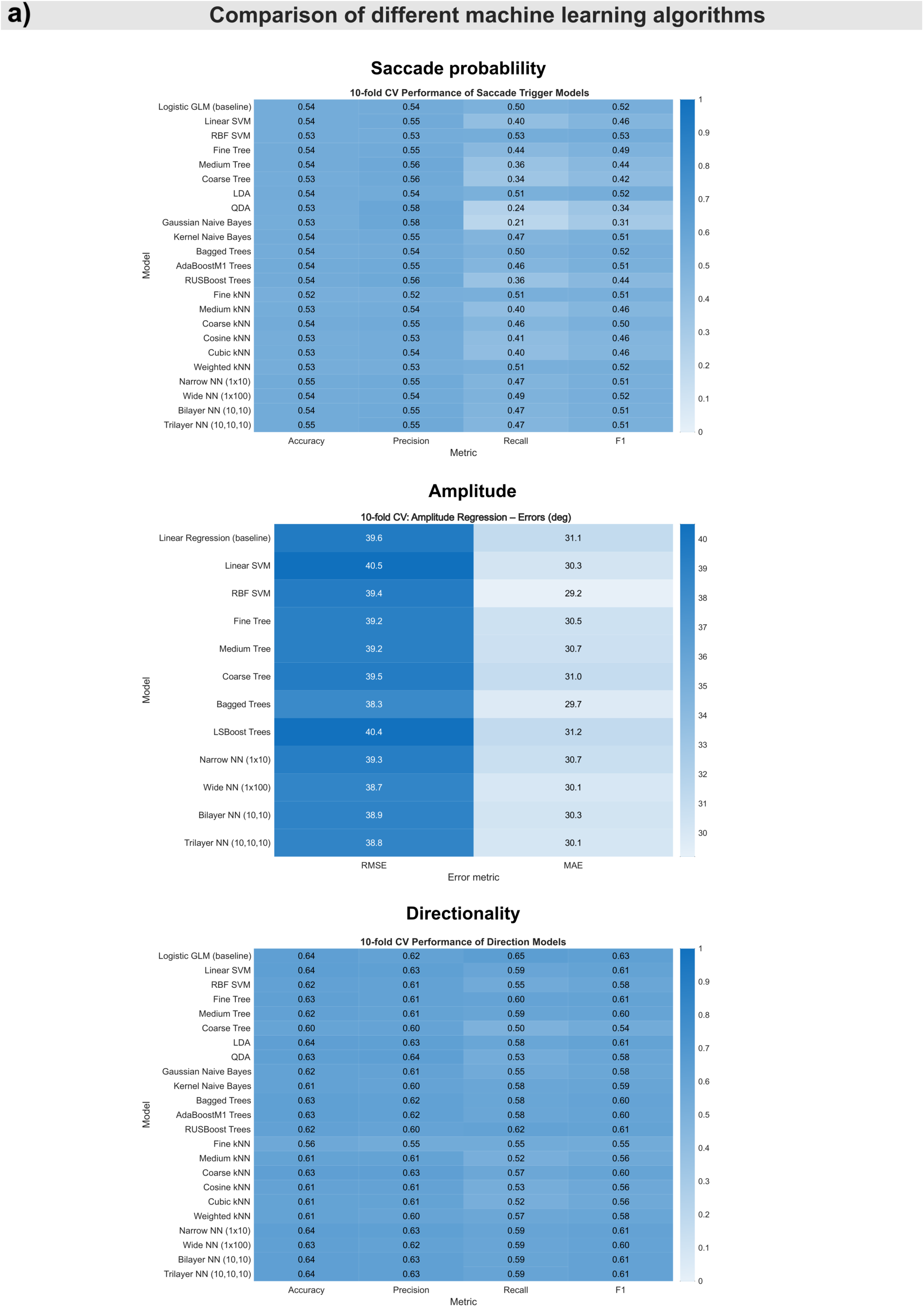
Comparison of different machine learning algorithms for predicting saccade trigger probability, saccade amplitude, and saccade direction. Models were trained on 80% of detected saccades and tested on the remaining 20% of non-training saccade data using 10-fold cross-validation. Top: Saccade probability. Middle: Saccade amplitude. Bottom: saccade directionality. For the classification models, we calculated accuracy, precision, recall, and F1. For the regression models (amplitude), we calculated the RMSE and MAE to determine model performance. More sophisticated modeling approaches (e.g., tree algorithms and neural networks (NN) performed similarly to the linear/logistic regression models used throughout this paper, but were also more difficult to interpret.

**Extended Data Fig. 5:**
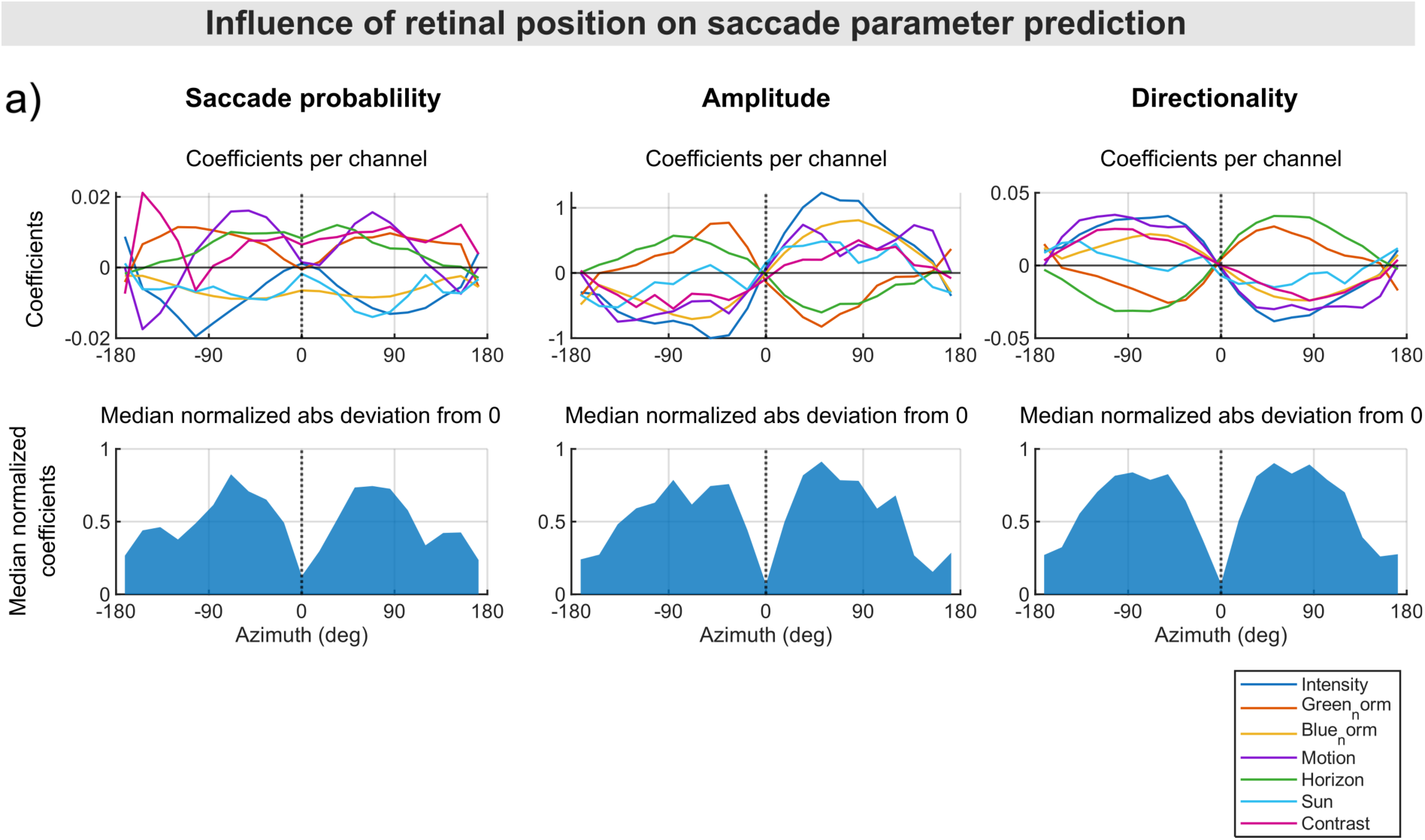
Influence of retinal position on saccade parameter prediction for saccade probability (left), saccade amplitude (middle), and saccade direction (right). 3 regression models (linear: amplitude, logistic: saccade trigger, directionality) were trained on the mean azimuthal visual input (downsampled into 21 bins) up to 500ms before saccades were performed. Top row: Coefficient estimates for the respective retinal positions are shown for each of the 7 analyzed visual parameters. Bottom row: Median normalized absolute deviation from 0 for all coefficients shown on top row for mapping the general influence of visual parameters on saccade parameters based on retinal position.

**Extended Data Fig. 6:**
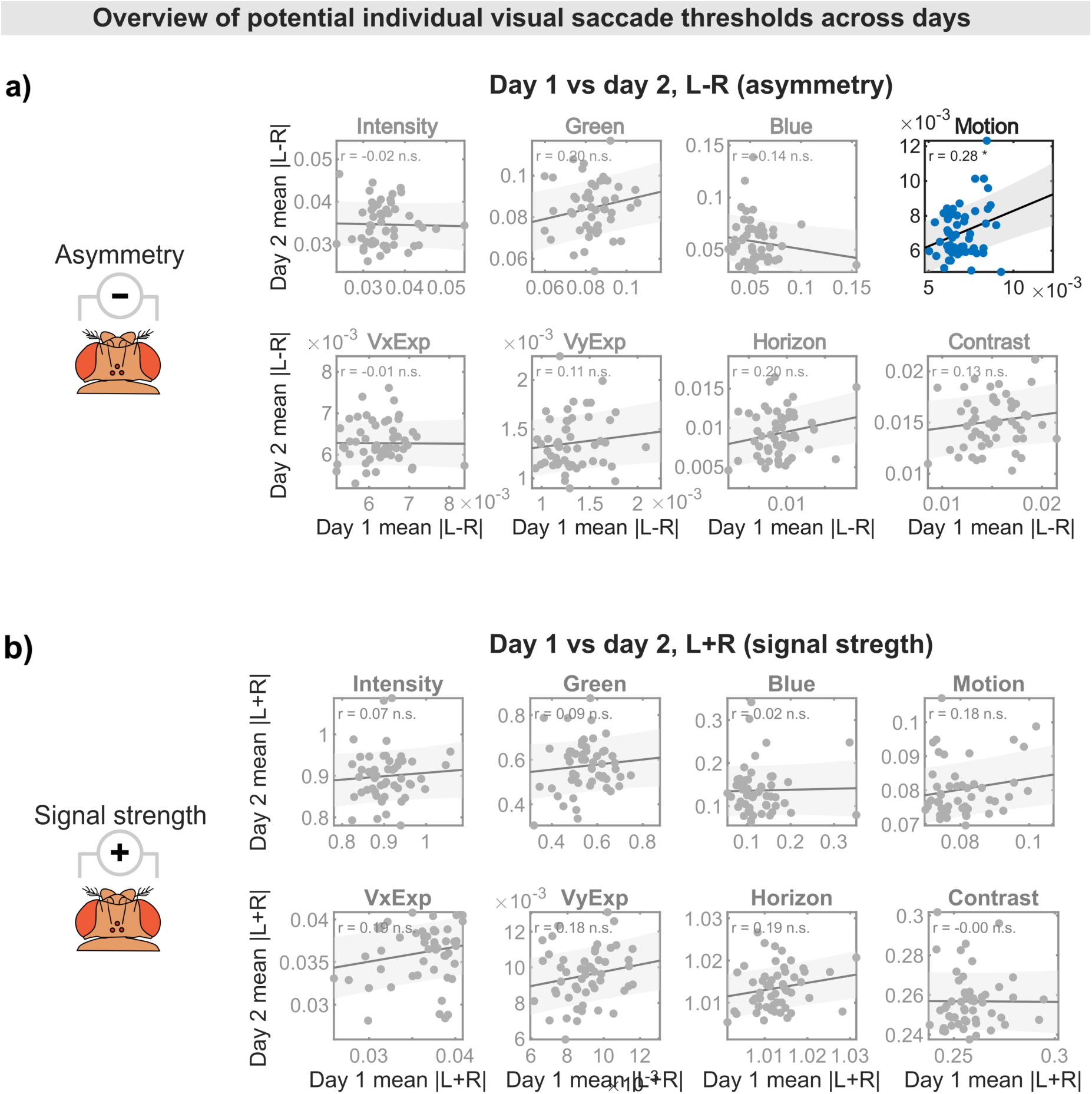
overview of potential individual visual saccade thresholds across days. **a)** Day-to-day correlation of Individual mean absolute visual input asymmetry (left minus right in the frontal visual field) experienced by flies up to 500ms before saccade initiation for 8 analyzed visual parameters. The individual absolute motion asymmetry levels after which flies perform saccades correlate significantly across days. Not significant correlations were shaded grey. **b)** Same as a) but for proposed left plus right frontal visual field computations. No significant day-to-day correlations were found. a-b) Correlations were quantified using Pearson’s correlation coefficient (r).

**Extended Data Fig. 7:**
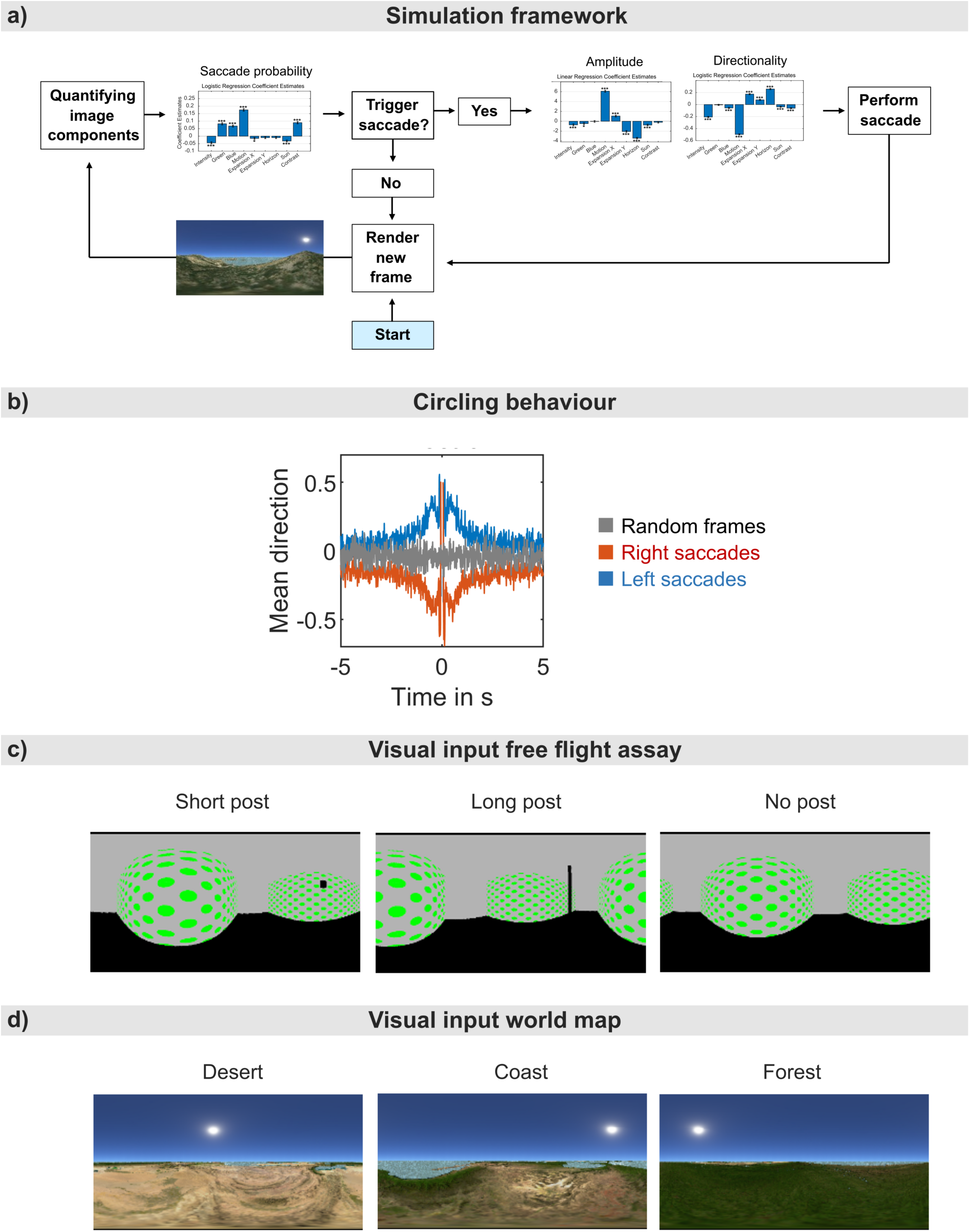
Simulating vision-based saccadic flight behavior. **a)** Simulation framework. The panoramic camera reads the first frame in Blender, which is loaded into MATLAB, and the image parameters are quantified. The saccade probability model (see Fig. 2 g) predicts, based on visual parameter input, whether to trigger a saccade. If not, the camera in Blender is moved forward along the previously determined direction, and a new frame is rendered and loaded into MATLAB. Once the saccade trigger model predicts a saccade, the visual parameters are sent to the other two models to predict saccade amplitude and direction. Rotational instructions for the panoramic camera are then sent to Blender, updating the camera position and azimuthal angle accordingly to simulate a saccade. After the saccade, a new frame is rendered, and the loop continues until 15 frames of virtual flight are simulated per fly with a sampling frequency of 80Hz. **b)** Circling behavior. All left (blue, 1) and right (red, –1) saccades (dataset from Fig. 2c) were aligned at timepoint 0s, and the mean saccade direction of saccades preceding/following these saccades was plotted from –5s to 5s. The probability of performing a saccade in the same direction as the previous saccade is temporarily increased. Grey: Mean saccade direction centered at randomly sampled frames. **c)** Modeling the free flight assay described by Maimon et al. (2008). 3 visual environments were modeled in 3D using Blender according to assay descriptions, containing a short black cylindrical post (left), a long black cylindrical post (middle), or no post (right) in the arena center. **d)** Simulating exploration behavior on a world map. 3 panoramic camera renders illustrating different virtual biotopes (left: desert, middle: coast, right: forest).

**Supplemental Video 1**

(Supplemental Video 1 – Virtual Reality.mp4)

Virtual reality setup overview. Left: Flight simulator with magneto-tether apparatus in the center. Middle: The camera stream is imported to MATLAB, and the fly’s heading is extracted and, in combination with a constant forward velocity, transformed to 2D coordinates. Right: Rendered panoramic camera image displayed on the LED matrix. Tracked 2D coordinates are applied to the camera’s position to simulate translational movement.

**Supplemental Video 2**

(Supplemental Video 2 – 2D Trajectories of Tested Flies.mp4)

2D trajectories of tested flies over a 15min test period (n=106). Right: zoomed-out view.

**Supplemental Video 3**

(Supplemental Video 3 – 2D Trajectories of Simulated Random Flight.mp4)

2D trajectories of simulated flies over a 15min test period (n=106). Right: zoomed-out view.

**Supplemental Video 4**

(Supplemental Video 4 – Time on Land.mp4)

Flies prefer flying over land. Left: masked island with positions of tested flies marked in red (n=106). Right: percentage of flies flying over land over the 15min test window. Blue trace: tested flies (n=106), grey trace: simulated flies performing random flight (n=106).

**Supplemental Video 5**

(Supplemental Video 5 – Explored Area.mp4)

Flies explore almost the whole island. Left: explored area over time (light blue, 2m exploration radius around each fly). Right: total percentage of the island explored over the 15min test window. Blue trace: tested flies (n=106), grey trace: simulated flies performing random flight (n=106).

**Supplemental Video 6**

(Supplemental Video 6 – Height Preference.mp4)

Flies prefer flying over high elevations. Left: height map of the island with flies marked in red (n=106). Right: median height of the fly population over the 15min test window. Blue trace: tested flies (n=106), grey trace: simulated flies performing random flight (n=106).

**Supplemental Video 7**

(Supplemental Video 7 – Vegetation Preference.mp4)

Flies prefer flying over vegetation. Left: vegetation (black) map of the island with flies marked in red (n=106). Right: mean vegetation preference of the fly population over the 15min test window. Blue trace: tested flies (n=106), grey trace: simulated flies performing random flight (n=106).

**Supplemental Video 8**

(Supplemental Video 8 – Brightness Preference.mp4)

Flies prefer flying over low intensity terrain. Left: brightness map of the island with flies marked in red (n=106). Right: mean brightness preference of the fly population over the 15min test window. Blue trace: tested flies (n=106), grey trace: simulated flies performing random flight (n=106).

**Supplemental Video 9**

(Supplemental Video 9 – Hue Preference.mp4)

Flies prefer flying over green terrain. Left: hue map of the island with flies marked in red (n=106). Right: mean brightness preference of the fly population over the 15min test window. Blue trace: tested flies (n=106), grey trace: simulated flies performing random flight (n=106).

**Supplemental Video 10**

(Supplemental Video 10 – Visual Feature Extraction.mp4)

Quantifying visual features. Top row: Panoramic image sequences were re-rendered, centered at the flies’ visual midline. Bottom row: Extracted visual feature channels and mean azimuthal signal strength.

**Supplemental Video 11**

(Supplemental Video 11 – Hilltopping.mp4)

Visually guided height preference optimizes chances of encountering mating partners through minimizing inter-fly distances during exploration. Left: Inter-fly distances in a naturalistic environment containing elevation data. Grey lines link each fly to all other flies of the population. n=21 (randomly selected from the fly population described in Fig. 1). Middle: same as left but for flies flying on a flat island without elevation data (see Fig. 2e, n=21). Right: Mean inter-fly distance over time for the fly populations exploring the island shown in the first two panels. Shaded areas indicate ±1 SD.

